# Colitis-induced visceral pain recruits central neurotensin neurons that modulate colonic sensitivity

**DOI:** 10.64898/2026.02.27.708254

**Authors:** Yu-Ting Cheng, Natalie MacKinnon-Booth, Yingfu Jiao, Jordan R. Robbins, Murillo Duarte-Silva, Perry E. Mitchell, Yuchu Liu, Omer Barkai, Keunjung Heo, Biyao Zhang, Bruna Lenfers Turnes, Meenakshi Rao, Clifford J. Woolf

## Abstract

Inflammatory bowel disease produces debilitating visceral pain that remains a major clinical challenge. Notably, many patients experience persistent pain even after the inflammation resolves, indicating a sustained sensitization of central neural circuits that drives enduring pain. The brainstem parabrachial nucleus integrates interoceptive signals from the gastrointestinal tract to elicit both pain perception and affective responses. Using activity-dependent mapping and an RNAscope assay, we identified a neurotensin (NT)-expressing neuronal population in the lateral PBN (PBN_L_) that is selectively activated during dextran sulfate sodium-induced colitis. *In vivo* neural activity recordings demonstrate that PBN_L_ NT neurons encode colon-derived nociceptive signals in an intensity-dependent manner. Silencing these neurons attenuates colonic reflexes evoked by luminal distension and normalizes aberrant gastrointestinal transit and nociceptive licking behavior in colitic mice. Pharmacological blockade of NT signaling alleviates colitis-associated hypersensitivity. These findings identify a central neural population that encodes visceral inflammation and regulates peripheral organ function, and pinpoints neurotensin as a promising therapeutic target to treat colitis-induced visceral pain.

## Introduction

Pain associated with inflammatory bowel disease (IBD) affects up to 10 million people worldwide and arises from the activation of nociceptor afferents by inflammatory responses mediated by a compromised epithelial lining of the gut barrier^1–3^. Clinical hallmarks of colitis include spontaneous abdominal pain, visceral hypersensitivity, and altered bowel function, such as diarrhea and dysmotility^4,5^. Current pharmacological treatments primarily target mucosal inflammation; however, tissue healing verified by endoscopy or histopathology, does not reliably produce symptomatic relief, with 30–50% of patients continuing to report persistent pain even after resolution of their inflammatory flare^6,7^. This suggests that a central neural sensitization may contribute to the maintenance of visceral pain independently of ongoing peripheral pathology^8–10^.

The brainstem parabrachial complex serves as a key integrative hub that relays interoceptive and exteroceptive signals from the periphery to forebrain regions to elicit pain and affective perception^11–15^. Within this complex, the lateral parabrachial nucleus (PBN_L_) primarily receives nociceptive input from the spinal dorsal horn via the anterolateral pathway and is engaged in acute and chronic pain states^16–18^. Genetic and circuit-level dissection of PBN_L_ neuronal populations, together with an analysis of their neuropeptide signaling has begun to reveal a powerful leverage mechanism for modulating pain and its associated emotional disturbances^19–22^. However, despite accumulating evidence implicating the role of PBN_L_ in somatosensory pain^23–26^, the identity of those neuronal ensembles that encode visceral nociceptive signals, and if and how these influence organ function and visceral pain-related behavior, are unclear.

We identified a population of neurotensin (NT)-expressing neurons within the PBN_L_ that is selectively activated during dextran sulfate sodium (DSS)-induced colitis and encodes colon-derived nociceptive information in an intensity-dependent manner. Silencing the PBN_L_ NT neurons abrogated the exaggerated colonic reflexes evoked by pressure distension, while blockade of neurotensin signaling normalized the diarrhea and pain-associated behaviors induced by the colitis. These findings provide evidence for a central neural modulation of peripheral visceral function and pinpoint PBN_L_ NT neurons as a mechanistically defined node linking visceral inflammatory pain and affective states, with promising implications for treating colitis-induced visceral pain.

## Results

### Activation of nociceptors by colitis drives spontaneous nocifensive behaviors

We modeled experimental colitis by administering 3% DSS into the drinking water of mice^27,28^ for 4 days and measured spontaneous pain-related behaviors together with abdominal allodynia for 2 weeks to evaluate the temporal profile of altered nociceptive behavioral responses (**Fig. 1a**). Colon inflammation was confirmed by an infiltration of CD45-positive immune cells into distal colon tissues 4 days after DSS exposure (D4) and 3 days after the DSS withdrawal (D7), which progressively resolved over 2 weeks (**Fig. 1b,c**). Additionally, we monitored a reduction of mouse body weight and water intake in DSS-treated WT mice compared to regular H_2_O drinking water controls, which gradually recovered on D14 (**Extended Data Fig. 1a,b**).

**Fig 1.**
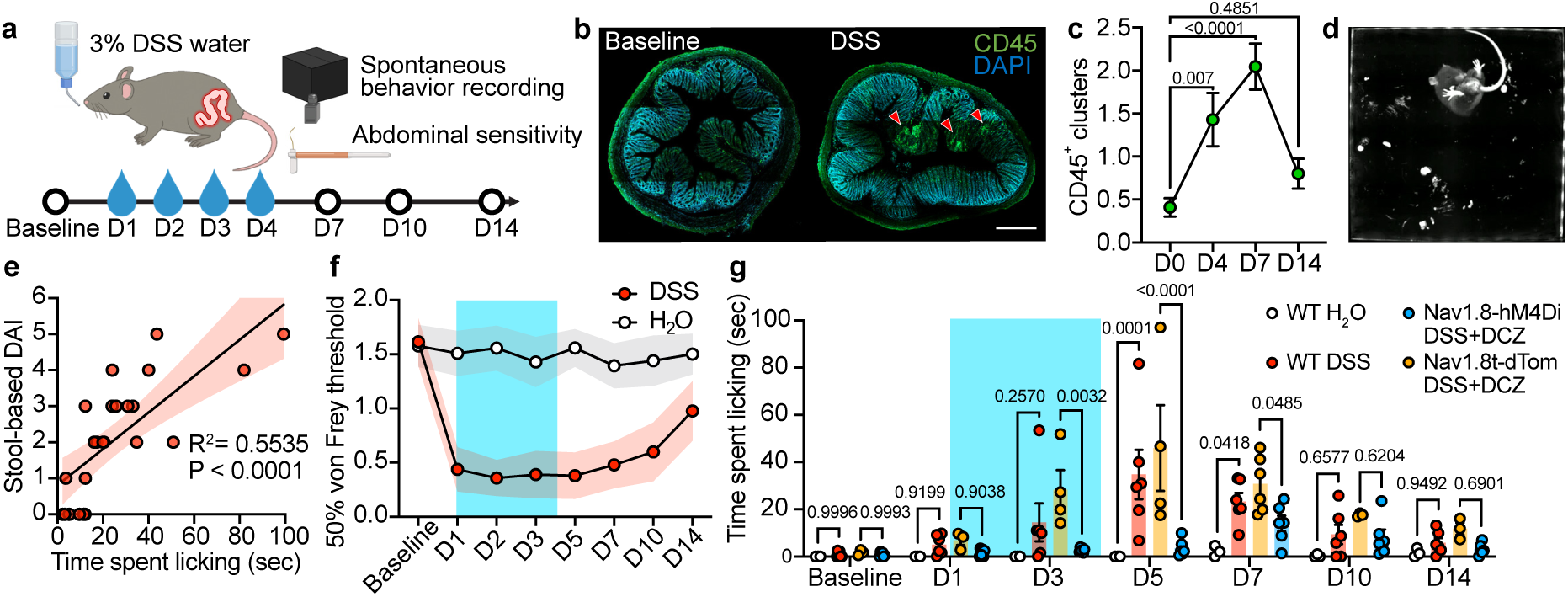
Nocifensive behaviors elicited in response to DSS-induced colitis. (a) Schematic of experimental timeline to evaluate spontaneous behavior and abdominal sensitivity in mice treated with 4-day of 3% dextran sodium sulfate (DSS) in their drinking water. (b) Histological verification of immune cell cluster (CD45^+^, green) invasion into the lamina propria of the mouse colon 3 days post 4-day DSS treatment (right) compared with baseline (left). Scale bar, 20 mm. (c) Quantification of CD45^+^ clusters at D0 (n=22 from 4 mice), D4 (n=14 from 3 mice), D7 (n=22 from 4 mice) and D14 (n=15 from 3 mice). One-way ANOVA with Dunnett’s multiple comparisons. (d) Example of a mouse with DSS-colitis engaged lower body licking behavior. (e) Linear regression analysis between the stool-based disease activity index and time spent licking over a 30-min recording session in mice exposed to DSS (n=25 mice). Light shaded areas represent 95% confidence band. (f) 50% cutaneous mechanical threshold of the mouse lower abdomen using an up-down von Frey test (H_2_O n=9 mice, DSS n=9 mice). Light blue shade: 3% DSS drinking water from D1-D4. (g) Automatic licking behavior quantified using a supervised machine learning classifier in WT mice with H_2_O (n=3), WT mice with 4-day DSS (n=6), Nav1.8-tdTom mice with 4-day DSS (n=3-6) and Nav1.8-hM4Di mice with 4-day DSS (n=6). In two Nav1.8 groups, the chemogenetic agonist deschloroclozapine (DCZ, 0.1 mg/kg) was injected i.p. at D1, 2, 3 and D4, once per day (two-way ANOVA with Tukey’s multiple comparisons).

We implemented a supervised machine learning algorithm to automatically screen for pain-related behavior signatures relevant to the DSS-induced colitis^29,30^. This analysis revealed a significant increase in “lower body licking” encompassing abdominal and peri-anal licking during the active inflammation, when compared with mice receiving regular H_2_O drinking water (**Fig. 1d**). A licking bout was defined as contact between the snout and the abdominal or peri-anal region sustained for at least two consecutive video frames. To interpret model features underlying licking classification, we analyzed the 10 most informative features using SHAP (Shapley additive explanations). Spatial features contributed most strongly to the model performance, with the distance between the snout/neck and the tail base ranked as the top feature, reflecting close proximity of the head to the lower body during licking behavior (**Extended Data Fig. 2a**). Model performance improved with increasing training data, as shown by the F1-score learning curve, and reached a plateau near 0.8 after approximately 6,000 labeled frames (**Extended Data Fig. 2b**). To further optimize classification performance, we evaluated precision, recall, and F1-score across a range of discrimination thresholds on the validation dataset. This analysis identified an optimal operating threshold centered around ∼0.6, which was used for subsequent behavioral quantification (**Extended Data Fig. 2c**).

A correlation between time spent licking and the stool-based disease activity index (DAI), a score calculated as the sum of two components: stool consistency and fecal blood^28^, demonstrated a positive linear relationship in the DSS mice (**Fig. 1e**), suggesting that lower body licking reflects the severity of colitis in mice. To measure colitis-induced mechanical allodynia, which is analogous to the feature of abdominal tenderness on clinical examination, we performed an up-down von Frey assay on the skin of the lower abdomen to calculate the 50% withdrawal threshold. We observed a profound decrease in the abdominal response threshold during the DSS water treatment, which gradually recovered after DSS removal to thresholds found in regular H_2_O regimen control mice (**Fig. 1f**). To examine the functional relevance of the licking behavior during the 4-day DSS water regimen to pain, we used Nav1.8-Cre mice that express either an inhibitory hM4Di receptor for chemogenetic inhibition, or tdTomato as a control, in dorsal root ganglion nociceptor neurons that trigger nocifensive reflexes in response to noxious stimuli (**Extended Data Fig. 3**). Mice where Nav1.8-lineage nociceptors were temporarily silenced by activating hM4Di on receiving 0.1 mg/kg of deschloroclozapine (DCZ)^31^ once per day over the 4 days of DSS exposure, displayed a marked reduction of nociceptive licking behavior and an attenuation of body weight loss (**Fig. 1g and Extended Data Fig. 1a**). This analgesic-like effect persisted on D5 after the cessation of Nav1.8 nociceptor silencing, but subsided by D7, when the behavior was indiscernible between Nav1.8-tdTom and WT controls with DSS treatment. This reveals that the behavioral licking signature tracks colitis severity and is triggered by Nav1.8-lineage nociceptors.

In parallel, we used an acute, mechanically induced visceral pain model by introducing graded mechanical distension in the mouse distal colon (**Extended Data Fig. 4a,b**). To produce an acute noxious colonic stimulus, a balloon was inserted and connected via a bifurcated tube to an air-pressurized syringe and a manometer to enable controlled distension and pressure monitoring. Acute colon distension at a pressure of 40 and 80 mmHg elicited pressure intensity-dependent spontaneous behaviors, including abdominal licking, abdominal squashing and hunching (**Extended Data Fig. 4c**), which were effectively diminished following Nav1.8-nociceptor chemogenetic silencing by exposure to DCZ (**Extended Data Fig. 4d-g**).

Additionally, we performed a cFOS analysis in the PBN_L_ of mice that received a colon balloon placement either as a sham, or with a pressure distension of 80 mmHg to identify central visceral pain-activated neurons. We found a substantial increase of cFOS-positive neurons in PBN_L_ following 30-min of stimuli and that the signal was significantly reduced by chemogenetic silencing of the Nav1.8-lineage neurons (**Extended Data Fig. 4h,i**), suggesting that peripheral Nav1.8^+^ nociceptors are the active driver of visceral pain hypersensitivity and trigger transmission of visceral pain signals into the central nervous system (CNS) to generate adaptive behavioral responses.

### Colitis drives a PBN_L_ neural activation mediated by peripheral nociceptors and inflammation

Neurons in the mouse parabrachial nucleus are molecularly diverse with multiple subsets of cell types that reside within distinct spatial niches^20–22^. We performed activity-dependent mapping in PBN neurons using cFOS expression to determine the neural activation pattern along the PBN_L_, a critical hub that receives multiple afferent inputs from the periphery and projects to the brain to generate sensory and affective responses. Following the 4-day DSS colitis induction paradigm we observed a significant increase in cFOS-positive signals in the PBN_L_ region compared to regular H_2_O controls (**Fig. 2a-c**). To verify whether this activation in the PBN_L_ is driven by primary sensory neurons, we utilized the Nav1.8-hM4Di mice for a temporary chemogenetic silencing of nociceptors, and TRPV1-DTA mice for a selective diphtheria toxin-based permanent ablation (DTA) of peripheral nociceptors. We observed a considerable reduction of the colitis-induced cFOS activation following either temporary or permanent silencing of Nav1.8 and TRPV1-lineage neurons, respectively (**Fig. 2a-f**).

**Fig 2.**
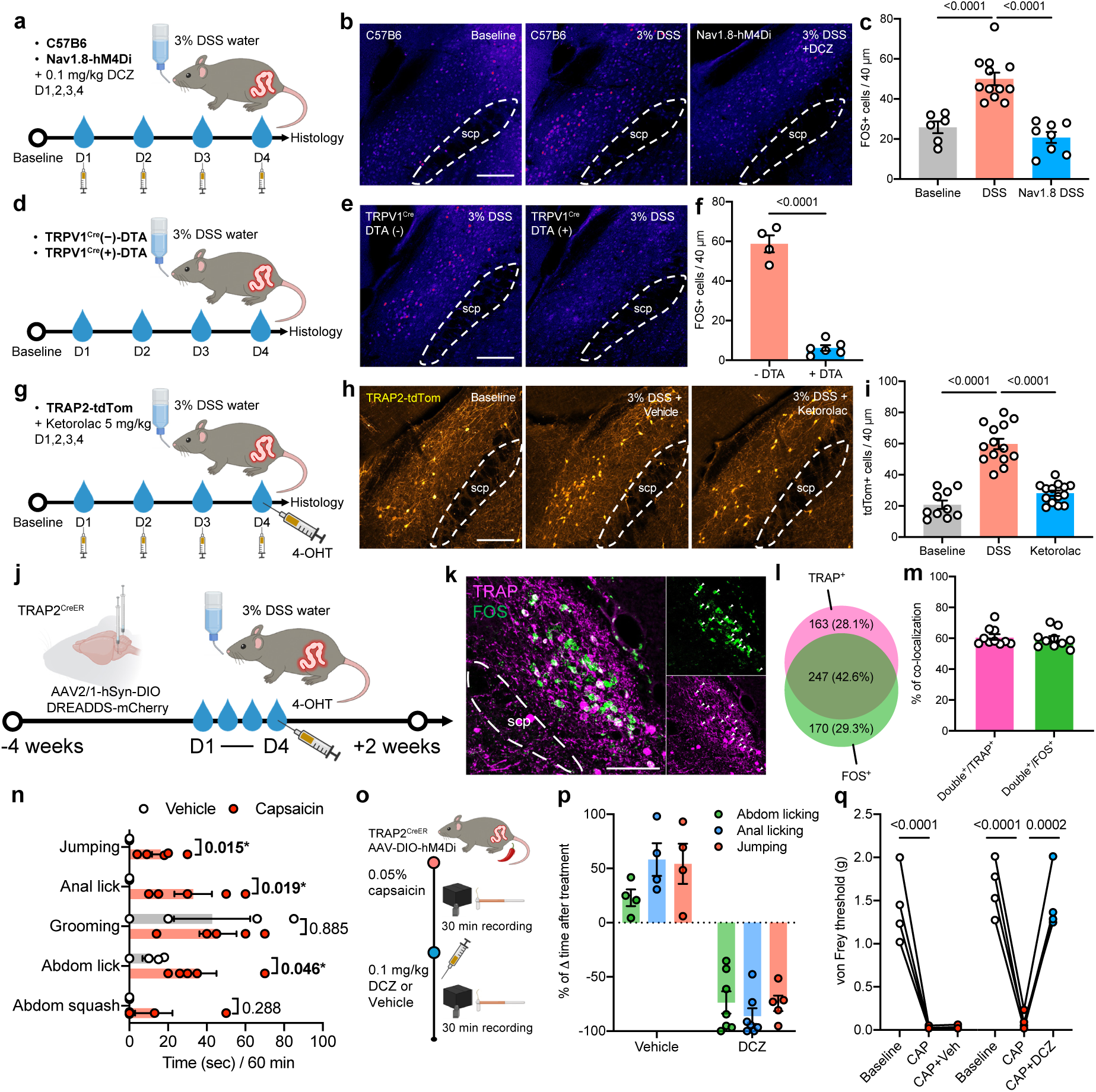
PBN^colitis^ neurons modulate behavioral responses to nociception in the colon. (a) Experimental scheme to measure FOS activation in the lateral parabrachial nucleus (PBN_L_) in WT and Nav1.8-hM4Di mice with 3% DSS drinking water. (b) Histological images of FOS antibody staining in PBN_L_ region at baseline, with exposure to 3% DSS and after Nav1.8 chemogenetic silencing. scp, superior cerebellar peduncle. Scale bar, 200 µm. (c) Quantification of FOS-positive signals (baseline n=3 mice, DSS n=6 mice, Nav1.8-hM4Di n=4 mice, one-way ANOVA with Tukey’s multiple comparisons). (d) Experimental scheme to measure PBN_L_ FOS activation in TRPV1^Cre^(-)-DTA and TRPV1^Cre^(+)-DTA mice with DSS. (e) Histological images of FOS staining in PBN_L_ region of TRPV1^Cre^(-)-DTA and TRPV1^Cre^(+)-DTA mice with DSS. Scale bar, 200 µm. (f) Quantification of FOS-positive neurons (TRPV1^Cre^(-) n=2 mice, TRPV1^Cre^(+) n=3 mice, two-tailed unpaired t-test). (g) Experimental scheme to quantify PBN_L_ FOS activation using TRAP2-tdTom mice under DSS treatment with i.p. injections of vehicle or 5 mg/kg ketorolac from D1-D4. (h) Histological images of tdTom^+^ cells in PBN_L_ region at baseline, 3% DSS with vehicle injections and 3% DSS with ketorolac injections. Scale bar, 200 µm. (i) Quantification of tdTom^+^ cells (baseline n=5 mice, DSS+vehicle n=7 mice, DSS+ketorolac n=7 mice, one-way ANOVA with Tukey’s multiple comparisons). (j) Experimental strategy to functionally label DSS colitis-activated parabrachial neurons (PBN^colitis^) using TRAP2-Cre^ER^ mice injected with Cre-dependent AAV vectors encoding chemogenetic actuators (hM4Di or hM3Dq) bilaterally into PBN_L_ followed by an i.p. injection of 4-hydroxytamoxifen (4-OHT, 10 mg/kg) to drive expression of FOS-positive cells after 4-day of 3% DSS treatment. (k) Histological verification of signal overlap between TRAPed PBN^colitis^ cells (magenta) and FOS RNAscope (green) under DSS colitis. Scale bar, 200 µm. White arrowheads indicate overlapping signals. (l) Pie chart depicting overlap between TRAP^+^ and FOS^+^ neurons. (m) Quantification of signal overlaps in TRAP^+^ versus FOS^+^ population (n=5 mice). (n) Quantification of spontaneous pain-related behaviors in response to an intracolonic 0.05% capsaicin infusion over a 60-min recording session (vehicle n=4 mice, capsaicin n=5 mice, multiple unpaired t-tests using two-stage step-up method (Benjamini, Krieger and Yekutieli)). (o) Experimental timeline to evaluate the effect of silencing PBN^colitis^ neurons on capsaicin-induced spontaneous and evoked nocifensive responses. (p) Percentage time difference of capsaicin-induced abdominal licking, peri-anal licking and jumping after vehicle or 0.1 mg/kg DCZ to silence PBN^colitis^ activity in a 30-min recording session (vehicle n=4 mice, DCZ n=5-7 mice). (q) Cutaneous threshold of mouse lower abdomen using up-down von Frey test (vehicle group n=4 mice, DCZ group n=4 mice, one-way ANOVA with Tukey’s multiple comparisons).

To selectively capture neurons activated during DSS-induced colitis, we employed an activity-dependent genetic labeling strategy by crossing TRAP2 mice with Rosa-lxl-tdTomato reporter mice. TRAP2-tdTom mice were subjected to 4-day DSS treatment followed by an administration of 4-hydroxytamoxifen (4-OHT). This approach enabled permanent labeling of FOS-expressing neurons that were active during the defined colitis window, allowing identification and analysis of DSS colitis-activated neuronal populations (**Fig. 2g**). We observed a comparable increase in the tdTomato-positive signal relative to baseline as the cFOS staining, and this increased FOS-driven activation was noticeably diminished in mice that received ketorolac injections, a nonsteroidal anti-inflammatory drug, to dampen the DSS-induced local inflammation in the colon (**Fig. 2h-i**). These results indicate that the central activation of PBN_L_ neurons is driven by peripheral inflammation and nociceptor inputs.

### PBN^colitis^ neurons tune abdominal sensitivity

Next, we sought to probe what function the activated PBN_L_ ensemble may have in mediating colitis pain and its associated affective phenotypes. To gain functional access to the cFOS-driven population, we utilized TRAP2 mice to functionally label the colitis-activated neural population. The TRAP2 mice first received a stereotaxic injection of AAVs encoded with Cre-dependent DREADDS bilaterally into PBN_L_. Colitis-activated PBN_L_ (PBN^colitis^) neurons were tagged by treating the mice with DSS water followed by an injection of 4-OHT to induce iCre recombination specifically in the FOS-expressing population (**Fig. 2j**). Quantification of the signal overlap between TRAPed and FOS-positive cells induced by DSS colitis revealed a ∼60% TRAPing efficiency for the TRAPed and FOS-positive population (**Fig. 2k-m**). We then asked whether PBN^colitis^ neuronal activity is sufficient to modulate basal mechanical thresholds in the abdominal region, a measure of the referred cutaneous hypersensitivity associated with visceral pain (**Extended Data Fig. 5a**). We found that chemogenetic silencing (hM4Di) or activation (hM3Dq) of the PBN^colitis^ neurons bidirectionally tuned abdominal mechanical sensitivity across a range of von Frey forces following DCZ administration when compared with vehicle controls (**Extended Data Fig. 5b,d**).

When the same cohort of mice was assessed for basal hindpaw mechanical sensitivity, neither silencing nor activation of PBN^colitis^ neurons produced any observable change in the paw withdrawal responses (**Extended Data Fig. 5c,e**). Hindpaw zymosan-activated PBN_L_ (PBN^Zymosan^) neurons modified the paw withdrawal response to von Frey stimuli (**Extended Data Fig. 7a,b**) and thermal sensitivity and preference across a temperature gradient (**Extended Data Fig. 7d-f**), but displayed a negligible modulatory effect on abdominal sensitivity (**Extended Data Fig. 7c**). These results support a modality-specific role of the PBNL in visceral versus somatic nociceptive processing.

### PBN^colitis^ neurons gate behavioral responses towards visceral nociception

To evaluate the impact of PBN^colitis^ neural activity in modulating colon pain-associated behaviors, TRAP2 mice with Cre-dependent inhibitory hM4Di AAVs received a colonic infusion of 0.05% capsaicin to activate TRPV1-lineage nociceptors in the distal colon to initiate pain (**Fig. 2o**). We identified several distinct spontaneous behaviors, including lower body licking, jumping and adnominal squashing immediately following the low-dose colonic capsaicin exposure compared to vehicle treated controls (**Fig. 2n**). These spontaneous behavior bouts were effectively reduced by silencing of PBN^colitis^ neurons with DCZ, as opposed to the increase in behavior bouts in vehicle treated mice (**Fig. 2p**). In parallel, we identified the development of tactile allodynia over the abdominal region in the colonic capsaicin-treated mice and that mechanical threshold returned close to baseline in response to PBN^colitis^ neural silencing (**Fig. 2q**).

### PBN^colitis^ neurons promote aversive states

To assess whether PBN^colitis^ neurons influence affective and motivational valences we implemented a three-chamber task where mice were allowed to freely explore a completely shielded chamber (“hidden zone”) or an open chamber (“open zone”). We then performed a conditioning paradigm by pairing the injection of DCZ in mice in the hidden zone, and vehicle in the open zone, in a set of mice that had previously received a stereotaxic delivery of Cre-dependent hM3Dq AAV for chemogenetic activation, or mCherry AAV as controls (**Extended Data Fig. 6a**). Real-time tracking of the mouse trajectory demonstrated that PBN^colitis^ activation drove an aversive response, as revealed by a robust reduction of the time spent in the hidden zone when that chamber was paired with DCZ treatment during the conditioning sessions (**Extended Data Fig. 6b,c**). In parallel, we also conducted a two-bottle choice test to monitor motivational changes towards an appetitive incentive (**Extended Data Fig. 6d**) and found that a strong taste aversion developed in mice after PBN^colitis^ activation conditioning, as indicated by significantly reduced consumption of sucrose when paired with DCZ treatment, compared with the control group (**Extended Data Fig. 6e,f**).

### PBN_L_ neurotensin neurons encode visceral pain induced by colonic insults

Cell type-specific populations reside in different subregions of the PBN_L_ and form distinct projection patterns and circuit functions^12,21,32^. We divided colonic stimulation induced cFOS expression in the PBN_L_ region into two distinct subregions, the external lateral PBN (elPBN) and the dorsal lateral PBN (dlPBN), based on established anatomical boundary standards^12,20,21^. We identified enriched cFOS-positive signals within the elPBN region following DSS-induced colitis, which comprised ∼67.5% of cFOS-positive signals within the elPBN compared to ∼32.5% within the dlPBN (**Extended Data Fig. 8**), indicating preferential activation of neural ensembles within elPBN under the condition of colitis. We then focused on the elPBN subregion to identify the neuronal cell types recruited during colitis using an established spatial transcriptomic dataset^20,21^. The elPBN region primarily encompasses a *Calca*-expressing population along with other gene markers, including *Tac1*, *Penk*, *Crh and Nts*. To focus on those elPBN neurons that send projections to sensory and pain-regulating brain regions, we used *Crh* and *Nts* markers to evaluate signal overlap in our cFOS dataset. We collected PBN_L_ from mice that had either received 3% DSS drinking water for 4 days to induce colitis or 20 µL of 5 mg/mL zymosan into the left hindpaw for 6 hours to induce plantar skin inflammation^33^ (**Fig. 3a**). We determined the signal overlap between the two marker genes and *Fos* expression by performing RNAscope on serial sections of PBN_L_ (**Fig. 3b**). *Nts^+^* neurons displayed a ∼65% overlap with *Fos^+^* signals in neurons during DSS colitis, with only a ∼30% signal overlap with *Crh^+^*neurons, and a ∼25% overlap between *Nts^+^* and *Fos^+^*signals in neurons following hindpaw zymosan (**Fig. 3c**), suggesting a preferential role of neurotensin-expressing PBN_L_ neurons in encoding visceral over somatic pain.

**Fig 3.**
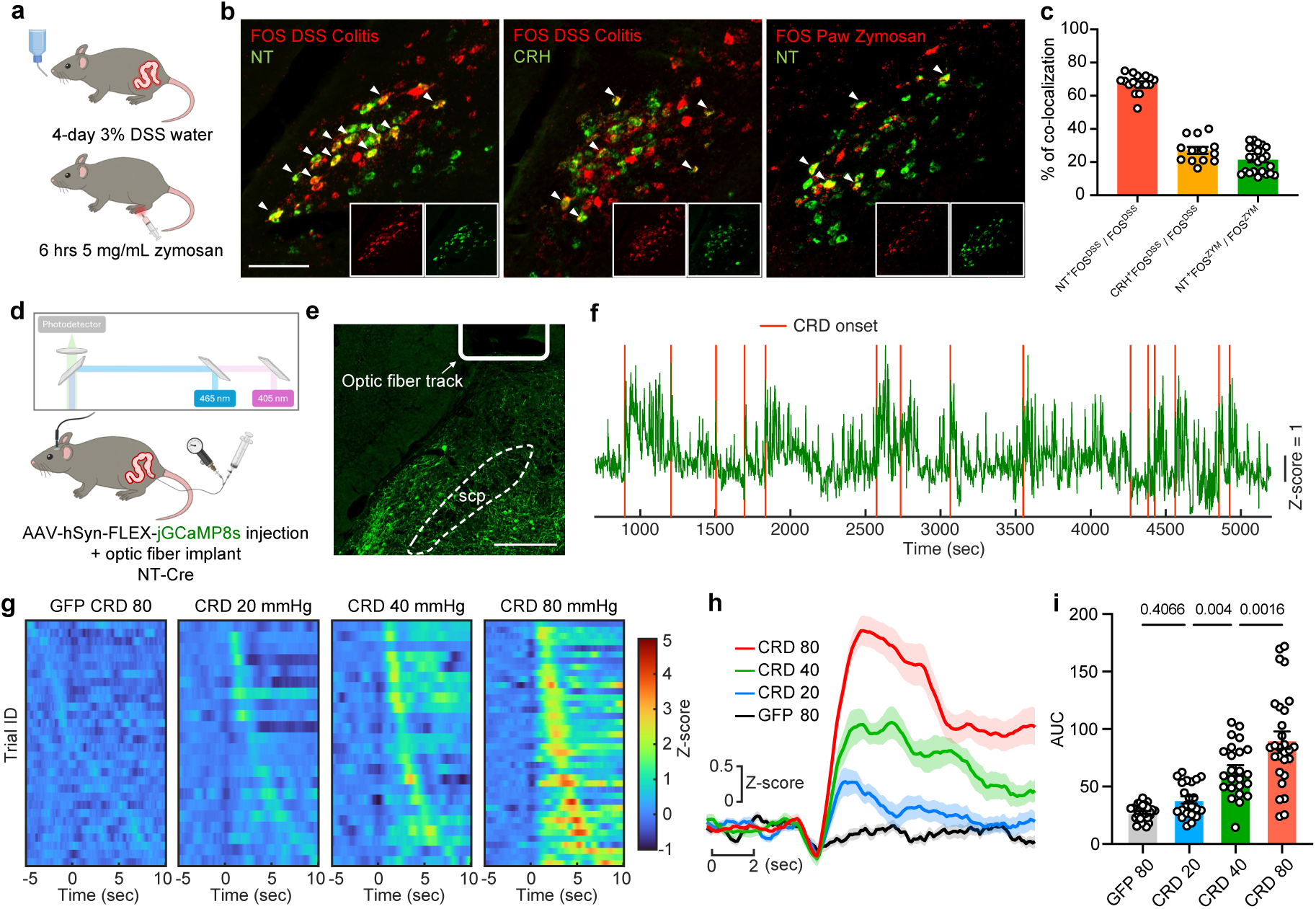
PBN_L_ neurotensin neurons encode colon-derived nociceptive signals. (a) Schematic of colitis-induced visceral pain in mice exposed to 4-day of 3% DSS drinking water (top) and somatosensory inflammatory pain induced by an injection of 5 mg/mL zymosan into plantar surface of the left hindpaw (bottom). (b) PBN_L_ sections with RNAscope detecting DSS colitis-activated FOS and neurotensin (NT) (left), DSS FOS and corticotropin-releasing hormone (CRH) (middle), and contralateral PBN_L_ sections with left hindpaw zymosan-activated FOS and NT (right). Scale bar, 200 µm. (c) Quantification of signal overlap between FOS DSS and NT (red bar), FOS DSS and CRH (orange bar), FOS Zymosan and NT marker (green bar). N = 17 sections from 4 mice, 13 sections from 4 mice and 24 sections from 5 mice. (d) Schematic of fiber photometric recordings of neural activity in PBN_L_ NT neurons evoked by intracolonic balloon distension. (e) Histological verification of AAV-transfected jGCaMP8s cells and fiber implant track in PBN of an NT-Cre mouse. Scale bar, 300 µm. (f) Example z-scored ΔF/F of GCaMP8s signal in NT neurons in response to colorectal distension (CRD) of 80 mmHg. (g) Heatmap responses of PBN_L_ NT neural activity to 10-sec CRD at 20, 40, 80 mmHg and GFP controls in response to CRD at 80 mmHg (GFP n=30 trials from 3 mice, CRD 20 n=22, CRD 40 n=24, CRD 80 n=40 trials from 5 mice). (h) CRD-triggered average z-scored ΔF/F of NT neurons. Light shaded areas represent s.e.m. (i) Area under curve (AUC) measured using average signal during the 5 sec prior to CRD onset as baseline, from (h). One-way ANOVA with Tukey’s multiple comparisons.

Using a neurotensin genetic driver of Cre mice (NT-Cre), we next sought to delineate the *in vivo* calcium dynamics of the PBN_L_ NT neurons using fiber photometry in response to the visceral pain elicited by noxious distension with a pressurized colon balloon (**Fig. 3d-f**). We inserted a balloon with a bifurcated ending to an air-pressured syringe and a manometer to monitor balloon distension pressure within the colon simultaneously (**Fig. 3d**) and introduced pressures of 20, 40 and 80 mmHg into the balloon to mechanically distend the colonic lumen with a 5-sec inflation to the designated pressure and a 5-sec deflation back to the baseline pressure. We observed an encoding of the PBN_L_ NT neural population to colonic distension in a pressure intensity-dependent manner (**Fig. 3g-i**), providing direct evidence for their role in the central processing of visceral afferent insults.

### PBN_L_ neurotensin neurons control visceral reflex and behavioral responses to colonic insults

We next sought to determine whether central NT-positive neurons play a functional role in colitis-induced symptoms. We utilized two different AAV-based silencing strategies to explore the impact of reduced PBN_L_ NT activity on the aberrant behavior and altered GI function triggered by DSS-colitis. We injected an AAV vector that encoded (1) Cre-dependent hM4Di (2) Cre-dependent TeLC (tetanus toxin light chain) or (3) Cre-dependent GFP into the bilateral PBN_L_ of NT-Cre mice and evaluated behavioral outcomes over a 4-day DSS treatment regimen (**Fig. 4a-c**). Temporary silencing elicited by M4 muscarinic receptor binding to the DREADD agonist versus long-term silencing mediated by a tetanus toxin-based synaptic blockade were implemented to compare their analgesic profiles on colitis-induced licking behavior (**Fig. 4d**). We found a robust attenuation of nociceptive licking behavior in mice that received 0.1 mg/kg of DCZ (once per day for 4 days) to silence PBN_L_ NT activity during DSS treatment, and that this analgesic effect persisted beyond the chemogenetic silencing period when compared to the GFP control group (**Fig. 4e**). Long-term silencing of NT release in the PBN_L_ by tetanus toxin substantially reduced the colitis-induced licking behavior with only a marginal increase in licking bouts at D7 (**Fig. 4e**).

**Fig 4.**
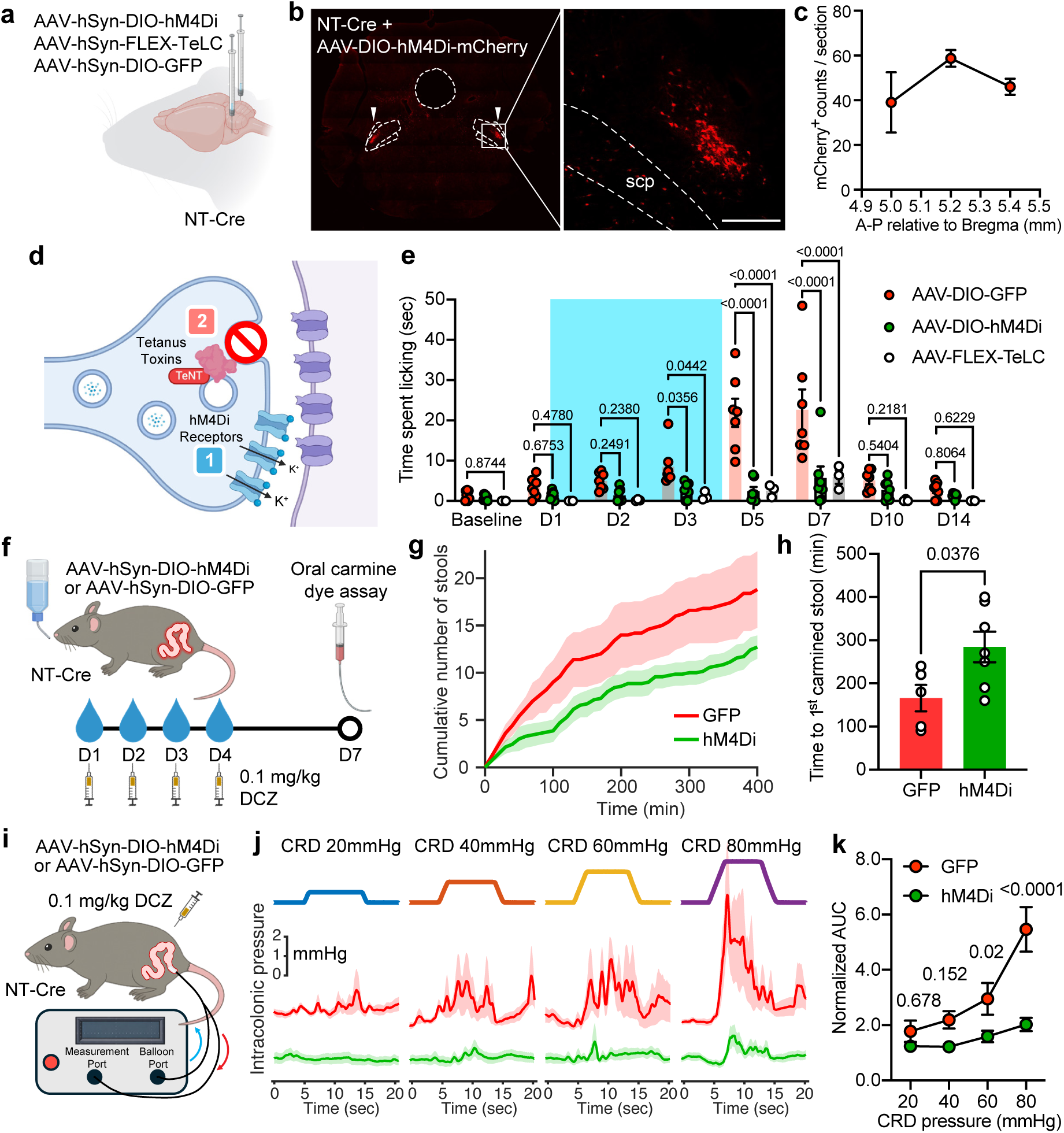
PBN_L_ neurotensin neurons control behavior and visceral sensitivity induced by colonic insults. (a) Schematic of two different silencing (hM4Di: modified human M4 muscarinic receptor for chemogenetic silencing, TeLC: tetanus toxin light chain-mediated synaptic impairment) and a control AAV GFP injection to measure causal function of PBN_L_ NT neurons. (b) Histological verification of Cre-dependent AAV injections into bilateral PBN_L_ of an NT-Cre mouse, scp, superior cerebellar peduncle. Scale bar, 200 µm. (c) Quantification of mCherry^+^ cells in the PBN_L_ at A-P axis 5.0, 5.2 and 5.4 mm relative to bregma (n=3 mice). (d) Diagram illustrating two different silencing strategies (1) virally mediated Gi-coupled muscarinic receptors induce an efflux of potassium ions for temporary neural hyperpolarization (2) virally mediated tetanus toxin light chain blocks a presynaptic fusion protein for long-term abrogation of neurotransmitter release. (e) Licking behavior quantified using a supervised machine-learning classifier in NT-Cre mice with bilateral injections of a Cre-dependent AAV that encodes GFP (n=7 mice), or hM4Di (n=7 mice) and TeLC (n=3 mice) in mice subjected to 4-day 3% DSS treatment. In both the GFP and hM4Di groups, the chemogenetic agonist (DCZ; 0.1 mg kg⁻¹, i.p.) was injected once daily on D1-D4. Two-way ANOVA with Dunnett’s multiple comparisons. (f) Experimental scheme to evaluate gastrointestinal transit using a carmine-red dye assay to measure latency of the first appearance of dye-colored stool following oral administration on D7. GFP and hM4Di mice underwent 4-day DSS treatment and received DCZ injection once per day from D1-D4. (g) Quantification of total stool number 30 min after an oral gavage of carmine dye followed by stool count every 10-min until 400 min (GFP n=5 mice, hM4Di n=7 mice). Light shaded areas represent s.e.m. (h) Quantification of latency to first carmine-dyed stool (GFP n=5 mice, hM4Di n=7 mice. Two-tailed unpaired t-test). (i) Schematic of colonic reflex response to graded distensions by an inserted balloon connected to a close-loop, calibrated barostat system. (j) CRD-triggered average intracolonic pressure responses to distension pressures of 20, 40, 60 and 80 mmHg in GFP (red, n=8 trials from 4 mice) or hM4Di (green, n=14 trials from 7 mice) mice. Light shaded areas represent s.e.m. (k) Quantification of normalized AUC (average of 5-sec AUC following CRD onset / average of 5-sec AUC prior to CRD onset) in response to CRD pressure at 20, 40, 60 and 80 mmHg in GFP (red, n=4 mice) and hM4Di (green, n=7 mice) group. Two-way ANOVA with Šídák’s multiple comparisons.

Licking bouts in mice are modulated either by a local inflammation-induced sensitization driven by Nav1.8-lineage nociceptors or through a body-axis projection that controls gastrointestinal (GI) physiology^34,35^. To test whether central NT neural activity modulates GI function, we delivered DCZ to NT-Cre mice that had received a Cre-dependent hM4Di or GFP AAV in the PBN_L_ during the DSS treatment, followed by an oral gavage of carmine dye at D7 to measure gastrointestinal transit time (**Fig. 4f**). In the NT-silenced group we found a normalization of the colitis-accelerated gut motility, as revealed by cumulative stool number over time and the latency to the first red fecal pellet expelled by the mice (**Fig. 4g,h**). We measured local colon sensitivity in response to a series of controlled distensions using a close-loop balloon barostat device in both the NT-silenced and control groups (**Fig. 4i**). Distension intensity-dependent changes of the colon reflex were present in control mice, with a large increase of the response when the distension pressure surpassed 60 mmHg. Importantly, these evoked responses were largely dampened in NT-silenced mice (**Fig. 4j,k**). We conclude that PBN_L_ NT neurons serve as a central hub to both sense visceral hypersensitivity and impact associated colonic reflex and behavioral outputs.

### PBN_L_ neurotensin neurons protect against colitis-induced pathology

To determine whether silencing PBN_L_ NT neurons attenuate the histopathological outcomes of DSS-induced inflammation, we administered a calibrated dose of DCZ into the mouse drinking water to achieve a sustained silencing of PBN_L_ NT neurons that matches the multi-day progression of the pathology (**Extended Data Fig. 10a**). Longitudinal assessment of the stool-based disease activity index (DAI) revealed that chronic NT-silencing significantly improved stool consistency and fecal bleeding, and reduced DAI compared to a non-silenced control group, as the disease progressed toward D7 (**Extended Data Fig. 10b**). Analysis of the colon revealed that silencing PBN_L_ NT neurons reduced neutrophil infiltration and preserved crypt integrity and overall colon length (**Extended Data Fig. 10c-e**). Fecal lipocalin-2, an inflammatory marker of mucosal immune activation, was significantly elevated at D7 under DSS colitis but was markedly reduced in the NT-silencing group (**Extended Data Fig. 10h**). These protective effects appeared to occur independently of overt systemic immune alterations, as evidenced by a lack of weight change in spleen and mesenteric lymph nodes (**Extended Data Fig. 10f,g**). These data indicate that PBN_L_ NT neurons can exert a protective influence against colitis-mediated pathology.

### PBN_L_ neurotensin neurons exert a restricted modulation of somatic inflammatory pain

We next investigated whether PBN_L_ NT neurons differentially modulate distinct peripheral sensory modalities. An intraplantar injection of zymosan was administered to induce inflammatory pain in the hindpaw. Somatic hypersensitivity was analyzed by measuring contact luminance of the hindpaws via a transillumination recording platform^30^. A reduction in contact luminance is a proxy for tactile allodynia as zymosan-induced inflammation reduces plantar surface contact (**Extended Data Fig. 9a,b**). Contrary to the effects observed in the colitis model, silencing of PBN_L_ NT neurons did not significantly attenuate the zymosan-induced hypersensitivity throughout the three-day post-injection period (**Extended Data Fig. 9c**).

### Blocking neurotensin signaling attenuates colitis-induced pain-related behavior

We next aimed to investigate the downstream brain targets that receive neural projections from the PBN_L_ NT neurons by conducting an AAV-based projection mapping, and identified several regions known to regulate pain, including the bed nucleus of the stria terminalis (BNST), central amygdala (CeA), ventral posteromedial (VM) thalamus and dorsolateral periaqueductal gray (dlPAG) (**Extended Data Fig. 11a**). We then evaluated whether blockade of downstream terminal NT release could provide any therapeutic relief of colitis-associated pain-related behaviors. To confirm a functional connectivity between colitis-activated PBN_L_ neurons and the terminal release of NT in the CeA, we delivered a Cre-dependent hM3Dq into the PBN_L_ region of TRAP2-Cre mice, followed by a 4-OHT injection during the 4-day DSS treatment to induce the labeling of PB^colitis^ by a FOS driver (**Extended Data Fig. 11b**). In the CeA, we delivered a G protein–coupled receptor activation–based neurotensin sensor (GRAB-NT1.0)^36^ followed by an optic fiber implant to monitor NT release dynamics *in vivo* (**Extended Data Fig. 11c**). We detected a robust surge in NT release in the CeA region after activation of PB^colitis^ by DCZ treatment, as compared to the baseline signal in the vehicle treatment group (**Extended Data Fig. 11d,e**), confirming NT signaling in the PBN-CeA pathway. To test whether blocking neurotensin receptor type 1 (NTSR1) signaling in the CeA region affects DSS-colitis behaviors, we implanted cannulas bilaterally in the CeA for selective NTSR1 antagonist, SR48692, infusions (0.3 μL/day at 20 nM) over 4-day DSS treatment (**Extended Data Fig. 11f**). This produced an effective attenuation of colitis-induced licking behavior bouts as well as a reduction in diarrhea, as evidenced by the normalization of the stool water content (**Extended Data Fig. 11g-i**). These results highlight CNS neurotensin signaling as a key modulator that calibrates colitis-induced visceral pain and associated gastrointestinal pathological changes.

## Discussion

Understanding the central neural modules that encode and regulate visceral sensation has been hindered by the diffuse and heterogeneous nature of the peripheral innervation of viscera and the absence of robust pain-related behavioral reflexes that allow direct functional assessment^4,6,37^. We implemented a supervised machine-learning framework that integrates pose estimation with body-part contact detection to extract subtle and disease-relevant behaviors. We identified a distinct lower body licking behavior that emerged selectively in mice with colitis. Notably, this behavioral signature closely tracked the temporal dynamics of the colonic inflammation and was functionally dependent on Nav1.8-lineage nociceptors.

By combining FOS-based activity mapping with TRAP to identify activation-defined neuronal populations, we found that DSS-induced colitis preferentially recruits neurons within the external lateral PBN, a region implicated in encoding aversive salience and the affective valuation of bodily threat^13,24,38^. Silencing this colitis-activated parabrachial ensemble (PBN^colitis^) markedly attenuated the behavioral responses to an acute intracolonic capsaicin challenge. In contrast to somatosensory pain, visceral pain is commonly characterized by emotional distress and is therefore thought to substantially engage limbic and affective circuits^39^. Consistent with this, we found that activation of PBN^colitis^ neurons robustly drives aversive behavior as assessed by both conditioned place and conditioned taste paradigms, indicating that this neural ensemble engaged during visceral inflammation promotes a negative valence.

The enrichment of FOS-positive neurons within the elPBN in response to colitis prompted us to define the molecular identity of the colitis-responsive cell types in this region. We discovered that neurotensin-expressing neurons exhibited a selective overlap with colitis-induced FOS activation, with minimal overlap with the FOS signals elicited by a zymosan-induced somatic inflammation in the hindpaw, supporting preferential engagement of the PBN_L_ NT population by visceral rather than somatic afferent inputs. Using *in vivo* fiber photometric recordings, we further demonstrated that PBN_L_ NT ensemble activity increased robustly in response to graded colorectal distension when intraluminal distension exceeded 40 mmHg. This threshold-dependent profile parallels the force-response characteristics observed in most colon-innervating dorsal root ganglion neuron subtypes^40,41^, supporting the notion that PBN_L_ NT neurons encode high-threshold visceral nociceptive signals.

PBN_L_ NT neurons constitute a distinct subset of the excitatory population within the elPBN region and partially overlap with *Calca*, *Oprm1* and *Tac1* markers, which have been implicated in pain modulation. We found that silencing the PBN_L_ NT population, using either chemogenetics, or toxin-directed impairment of synaptic transmission, generated a profound analgesic effect, as evidenced by a marked reduction in colitis-associated licking behavior. Notably, this analgesic effect was more sustained than that achieved by targeting Nav1.8-lineage nociceptors, indicating that PBN_L_ NT neurons operate at a higher-order integrative node in the visceral pain pathway. We reasoned that the improvement of the nocifensive behavior is contributed by a normalization of local hypersensitivity in the sensitized colon. Importantly, we identified a significant reduction in the colonic hyperreflex triggered by barostat distension with PBN_L_ NT silencing, particularly at pressures exceeding the 40 mmHg threshold associated with visceral pain. In addition, PBN_L_ NT silencing also restored gastrointestinal transit, emphasizing that this neuronal population integrates visceral sensory inputs to coordinate both pain-related behaviors and peripheral organ-level physiological responses.

Neurotensin has been implicated in the regulation of fundamental homeostatic processes, including feeding, social valence and pain^42–44^, and has also been characterized as a peripheral proinflammatory mediator, as evidenced by an elevated expression of its receptor, NTSR1 in the colon in experimental colitis models^45–47^. We hypothesized that central neurotensin signaling contributes to the visceral nociceptive and autonomic sequelae of colitis. Using AAV-based circuit tracing, we revealed that PBN_L_ NT neurons project to affective brain regions, with a particularly dense innervation of the central amygdala (CeA). Activation of this parabrachial source evoked a robust surge in NT release in the CeA, as detected by a neurotensin-based GRAB sensor. Importantly, pharmacological blockade of NTSR1 signaling within the CeA effectively ameliorated colitis-associated licking behavior and diarrhea, indicating that central NT signaling acts as a supraspinal modulator of disease severity, potentially through the amplification of those aversive and autonomic pathways that feed back onto peripheral visceral inflammatory and nociceptive processes^34,35,48^. By integrating machine learning-assisted behavioral decoding, molecular profiling, *in vivo* neural activity recording, and circuit-specific manipulations, our study provides evidence for a central circuit mediated modulation of peripheral visceral function. These findings offer a mechanistic framework for understanding those visceral disorders arising from dysregulated brain-body interactions, and pinpoint neurotensin in the CNS as a potential therapeutic target that can be leveraged to treat colitis-induced visceral pain.

## Materials and Methods

### Animals

All animal husbandry and procedures involving mice were conducted in strict accordance with guidelines set forth by the Boston Children’s Hospital Institutional Animal Use and Care Committee. Mice at 6-8 weeks of age with the strains of C57BL/6J (B6, JAX #000664), Fos^2A-iCreER^ (TRAP2, JAX #030323), Nav1.8-Cre (JAX #036564), NT-Cre (JAX #017525), R26-LSL-hM4Di/mCitrine (JAX #026219), Ai14(RCL-tdTomato)-D (JAX #007914), TRPV1-Cre (JAX #017769) and ROSA-DTA (JAX #009669) were obtained from the Jackson Laboratory. Adult mice of 8-12 weeks of age for both male and female mice of heterozygous alleles were used for this study. Induction of iCre-mediated recombination in the TRAP2 mice was performed by an i.p. injection of 0.1 mL of 10 mg/mL of 4-OHT dissolved in corn oil, with experiments beginning ∼2 weeks after the induction.

### Pharmacological reagents

Dextran sulfate sodium salt (MP Biomedicals #0216011080) was dissolved in regular drinking water at a concentration of 3% to induce colitis, 4-hydroxytamoxifen (Sigma-Aldrich #H6278) was prepared in corn oil and ethanol for the induction of Cre^ER^ translocation, deschloroclozapine (Hello Bio #HB9126) was prepared in saline solution as an agonist for DREADDs experiments, ketorolac (TargetMol #T1212) was prepared in saline, capsaicin (Tocris #0462) was dissolved in ethanol, tween80 and saline to induce acute nociceptive responses, zymosan A from saccharomyces cerevisiae (Sigma-Aldrich #Z4250) was prepared in saline solution for intraplantar injection to induce hindpaw inflammation and pain. SR48692 (Sigma-Aldrich #SML0278), a selective NTSR1 antagonist, was prepared at 20 nM in a solution of saline with 2% dimethylsulfoxide (DMSO). Carmine powder (Sigma-Aldrich #C1022) was prepared at 6% w/v with methylcellulose to measure gut transit time.

### Adeno-associated virus (AAV) reagents

For chemogenetic manipulation of PBN_L_ neurons the following AAV vectors were used: AAV2/1-hSyn-DIO-hM4Di-mCherry (BCH Viral Core; 3.51×10^13^ gc/mL) for silencing, AAV2/1-hSyn-DIO-hM3Dq-mCherry (BCH Viral Core; 2×10^13^ gc/mL) for activation and AAV2/1-hSyn-DIO-mCherry (BCH Viral Core; 3.48×10^13^ gc/mL) as fluorescent controls. For permanent silencing of PBN_L_ neurons AAV2/8-hSyn-FLEX-TeLC-GFP (BCH Viral Core; 1.56×10^13^ gc/mL) was used to block synaptic transmission. For fiber photometric recording AAV2/8-hSyn-FLEX-jGCaMP8s-WPRE (BCH Viral Core; 4.16×10^13^ gc/mL) was used to monitor neural calcium activity and AAV2/1-hSyn-FLEX-H2b-GFP (BCH Viral Core; 3×10^13^ gc/mL) for fluorescent controls. To detect neurotensin release AAV2/1-hSyn-GRAB-NT1.0 (Addgene plasmid #180006 from Y. Li, viral vector packaged by BCH Viral Core; 2.78×10^13^ gc/mL) was used.

### Stereotaxic injections and surgical procedures

#### Intracranial AAV vector injections

Intracranial injections were conducted with aseptic procedures on a digital stereotaxic platform for small animals. Mice were anesthetized with 4% isoflurane and maintained with ∼1.5% throughout the entire surgery on a feedback-controlled heating pad at a body temperature of ∼37°C. We injected AAVs using a 33 gauge, 5 μL microsyringe (Hamilton #65460-03) at a rate of 50 nL/min for 250 nL volume in one injection site. PBN_L_ was targeted at coordinates relative to bregma: AP= –5.20 mm, ML= ±1.43 mm, DV= –3.55 mm. CeA was targeted at coordinates: AP= 1.15 mm, ML= ±2.83 mm, DV= −4.17 mm. Mice were allowed to recover and waited for at least 3 weeks for the virus to express before experiments were conducted.

#### Optic fiber and cannula implantation

The optic fiber was placed ∼0.3 mm above the AAV injection site to monitor bulk calcium activity using fiber photometric recording. For pharmacological experiments, cannulae (22GA, PlasticsOne) were implanted bilaterally above the targeted brain region for repeated infusions of drug into the mouse brain. All implants were tightly anchored onto the skull and secured with Metabond (Parkell) and dental cement. Implanted mice were allowed to recover at least 2 weeks before experiments.

### Behavior assays

#### Spontaneous behavior recording

Mice were individually placed in a 18 × 18 × 15 cm (length x width x height) black acrylic chamber closed on all sides (BlackBox Bio) except for an NIR camera (Basler acA2000-50gmNIR GigE, Basler AG, Ahrensburg, Germany) was positioned 30 cm beneath the glass surface to track whole body movement from a bottom-up perspective^30^. Image brightness features were captured using LED light reflected changes in pixel intensity acquired at a frame rate of 25 Hz with 1000 × 1000 pixels dimensions. Mice were habituated in the recording chamber 1 hour per day for three days before the start of the experiment. Recordings were performed in both experiment and control groups on the same day for 30 min in each session during the animal’s light cycle.

#### Automated behavior screening

Lower body licking behavior was quantified using BAREfoot, a supervised machine learning framework that integrates pose estimation with light-based measurements of body-part pressure and spatial distance^29^. Training datasets for the lower body licking classifier were generated through manual annotation of video recordings by experienced scorers, after which new classifiers were trained on these labeled data. After testing and validating classifier performance, automated detection and quantification of lower body licking behaviors bouts was performed. Manual annotation of spontaneous behavior signatures, including peri-anal licking (repetitive licking around the anal port), abdomen licking (repetitive licking on a fixed region of the lower abdomen), abdomen squashing (contact with pressing of the lower abdomen against the floor), hunching (shortening the distance between fore and hindlimbs) and jumping (leaping off the floor with two hindlimbs), were labeled by three experienced independent investigators on a frame-by-frame level using a customized MATLAB scoring platform.

#### Paw luminance measurement

Hind paw coordinates were tracked using DeepLabCut^49^, and 23 × 23 squared pixels centered on these locations were extracted from the FTIR frames. Tactile allodynia was assessed by measuring paw luminance, defined as the average pixel brightness within each extracted ROI.

#### Abdominal and plantar von Frey assay

Mice were habituated in von Frey chambers on a meshed floor for three days (1 hour per day), including the day of the test, before the mechanical sensitivity was evaluated. To evaluate the reflexive sensitivity in response to mechanical stimuli, a logarithmically increasing set of 8 von Frey filaments (Stoelting Co, #58011) ranging from 0.02 to 4.0 grams were applied perpendicularly to the plantar hindpaw or lower abdomen with a sufficient force to gently bend the monofilament. We employed the up-down method to measure the 50% withdrawal threshold on the mice lower abdomen or plantar hindpaw^50^. The response frequency was measured as the number of positive rapid withdrawal of the lower abdominal region or the hindpaw in response to 10 von Frey stimulations at an interval of at least 30 sec between each stimulus. All the behavior tests were conducted in a blinded fashion to either the experimental vs control group, or the injection of treatment vs vehicle group.

#### Colorectal distension

Mice were briefly anesthetized with ∼1.5% of isoflurane to receive a lubricated polyethylene balloon insertion through the rectal port as previously described^51^. A colonic balloon with a diameter of ∼2 mm was gently placed at the distal colon segment ∼20 mm from the mouse anal verge and connected to a pressure-equalization tubing with a manometer (Cole Parmer) positioned at the bifurcation to simultaneously monitor distension pressure. A pressured tape was attached to the base of the tail to secure the placement of the colonic balloon for freely behavior recording. For each distension trial, the balloon was inflated to the designated pressure of 40 mmHg or 80 mm Hg within 5 seconds and a 5-sec deflation back to the baseline pressure for a total of 30-sec distension. Each behavioral recording consisted of alternation of 30-second distension and 30-second rest, continuously over a 30-minute recording session.

#### Conditioned motivation tests

To evaluate the impact of colitis-activated PBN_L_ neurons on motivational behaviors in mice, we conducted conditioned place motivation and conditioned taste motivation tasks. For conditioned place motivation, mice were habituated for 1 hour in a three-chamber apparatus separated with a central corridor that allowed free access to left or right compartment, including one shielded compartment with a meshed floor (hidden zone) and the other open compartment with a smooth acrylic floor (open zone). On day 1, mice were placed in the central corridor to freely explore both compartments for 30 min to determine the place preference by tracking the time spent across the entire apparatus using Ethovision XT (Noldus). On conditioning day 2 and 3, mice were confined to each compartment and paired with an i.p. injection of 0.1 mg/kg DCZ in the hidden zone, or vehicle in the open zone for 1 hour. The conditioning occurred twice a day (9 AM and 5 PM) with alternating compartments for a period of 2 days. On day 4, mice were re-introduced back into the central corridor to freely explore for 30 min to record the time spent in each compartment. For conditioned taste motivation, mice were single-housed and habituated to the home cages and water bottles for three days before the experiment. Prior to each day of conditioning, mice were restricted to water access overnight and the consumption amount was measured during the light cycle (9 AM to 5 PM). On day 1 and day 3, mice were allowed to have free access to regular drinking water in the home cage for 30 min, followed by an i.p. injection of vehicle, then returned to the home cage until 8 hours were timed to measure the consumption amount of solution. On day 2 and day 4, mice were given free access to a 10% sucrose solution in the water bottle for 30 min, followed by an i.p. injection of 0.1 mg/kg DCZ, then returned to the home cage until 8 hours were timed. On day 5, mice were given two bottles that contained regular water and 10% sucrose in each, for 8 hours to measure the taste preference. The preference index of sucrose over water during the two-bottle test on day 5: (amount of sucrose consumption) / (amount of water + sucrose consumption).

#### Thermal gradient ring (TGR)

Mice were habituated for 30 min in the thermal gradient ring (Ugo Basile #35550) with the platform temperature pre-set to 10°C on one side and 55°C on the opposite side at an increment of 5°C along the circular ring. During the test, each mouse was allowed to freely explore along the TGR setup for 1 hour and the occupancy duration as well as distance traveled in each temperature zone were recorded using the ANY MAZE software.

#### Hotplate assay

Thermal nociceptive thresholds were assessed using a hotplate assay. Mice were first acclimated to the device for 1 hour before the experiment. The hotplate (BIOSEB) was preset and maintained at a constant temperature of 52°C degree. Each mouse was placed within a transparent plexiglas cylinder on the heated surface, and the latency to exhibit the first nocifensive flinching behavior was recorded. A maximum cut-off time of 30 seconds was implemented to prevent plantar damage. Each mouse was tested in three trials with at least 10-minute between trials and the average latency is calculated.

#### Gastrointestinal (GI) transit assay

To assess total gastrointestinal transit time, mice were orally gavaged 300 μL of 6% carmine red powder dissolved in 0.5% methylcellulose as previously described method^52^. Mice were then placed individually in clean home cages to measure the number of expelled stools. The GI transit time was defined as the interval between oral gavage and the first observation of a carmine-colored fecal pellet with the observation conducted every 10 minutes to ensure temporal accuracy.

### Physiological recordings

#### In vivo fiber photometry recording in awake behaving mice

Optical fibers (400 μm core diameter, RWD) were implanted two weeks following Cre-dependent AAV delivery of GCaMP8s or GRAB-NT1.0. During implantation, real-time photometric signals were monitored to optimize fiber placement depth to maximize fluorescence signal intensity. Photometric recordings were acquired using a fiber photometry system (Doric Lenses) controlled by Doric Neuroscience Studio software. Excitation light at 405 nm (isosbestic, activity-independent control) and 465 nm (activity-dependent signal) was delivered, and emitted fluorescence signals were recorded at a sampling rate of 120 Hz. To enable temporal alignment with behavioral recordings, the fiber photometry system generated a synchronization light pulse every 90 s, which was captured concurrently by a behavior video camera operating at 30 Hz. Raw photometry signals were processed by averaging every 12 consecutive samples, yielding an effective sampling rate of 10 Hz. To directly compare neural activity and behavior response to stimulus onset, fiber photometry and video-derived behavioral signals were resampled onto a shared time base. Photometry data were exported to MATLAB (MathWorks) for further processing and analysis. Fluorescence signals were expressed as ΔF/F, calculated as ΔF/F = (F − F₀)/F₀, where F represents the 465 nm activity-dependent signal and F₀ denotes the fitted isosbestic control signal derived from the 405 nm channel. Z-scored ΔF/F values were used for synchronization and subsequent analyses.

#### Distension barostat recording

For assessment of colonic sensitivity, colorectal distension was performed using a close-loop, programmed balloon distension barostat (G&J Electronics Inc). Mice were briefly anesthetized with 3% isoflurane followed by an i.p. injection of either vehicle or DCZ. Mice were then loosely positioned in a cylindrical restrainer, and a lubricated flexible balloon catheter was gently guided into the anus and advanced into the distal colon. The catheter was secured by taping it to the tail and the restrainer was tightened to minimize movement. Mice were allowed to recover fully from anesthesia and habituate in the restrainer for 30 min prior to recording experiment. The barostat operated through a closed-loop feedback system and utilized an internal pressure transducer to monitor intracolonic pressure change in real-time. The distension protocol is consisted of balloon inflation to graded pressures of 20, 40, 60, and 80 mmHg for 10 sec per distension pressure, with a 3-min inter-stimulus interval between distensions. Each pressure series was repeated three times. The barostat control signal indicating balloon distension was synchronized with the intracolonic pressure recordings obtained from the probe sensor to define distension timing and pressure levels. All raw data were acquired and processed using LabChart software (ADInstruments). To isolate dynamic pressure fluctuations, the intracolonic pressure signal was processed offline using a high-pass Butterworth filter (cutoff frequency, 0.1 Hz) to remove low-frequency background drift. The root mean square (RMS) amplitude of the filtered signal was calculated within predefined analysis windows and stored as a derived signal. For each distension epoch, the area under curve (AUC) of the RMS signal was calculated relative to the pre-distension baseline and used as an integrated measure of intracolonic pressure activity. Identical filtering parameters and analysis windows were applied across all experimental sessions.

### Colon histopathology and stool composition analyses

#### Colitis severity evaluation

Colon tissues were prepared as swiss-rolls, fixed, and sectioned for Hematoxylin & Eosin (H&E) staining to assess the severity of colonic inflammation. Colitis severity was assessed using a semi-quantitative scoring system based on three parameters: crypt dropout, submucosal expansion, and neutrophil infiltration. Each parameter was scored on a scale from 0 to 3, where 0 represented absence of pathology and 3 indicated severe involvement. The scores from all three parameters were summed to yield a total histology score ranging from 0 to 9, with higher scores reflecting greater tissue damage and inflammatory severity. Disease activity was further assessed using a stool-based disease activity index (DAI), which was calculated as the sum of two components: stool consistency and fecal blood. Stool consistency was scored as follows: formed and soft=1, loose or semi-formed=2, diarrhea=3. Fecal blood was evaluated using a Hemoccult card (Beckman Coulter) to detect occult blood and scored as follows: occult blood=1, visible blood=2, rectal bleeding=3. For stool composition measurements, mice were transferred to clean cages and allowed free access to food and water. Stool output was continuously collected for 1 h. The total mass of fecal material collected during this period was recorded as the wet weight per mouse. Collected fecal pellets were then air-dried for 14 days until a stable mass was achieved, at which point the dry weight was recorded. Percentage of fecal water content = [(wet−dry)/wet]x100.

#### Fecal lipocalin-2 (LCN2/NGAL) quantification by ELISA

Fecal lipocalin-2 levels were quantified using a sandwich ELISA (DuoSet™ Mouse Lipocalin-2/NGAL, R&D Systems/Bio-Techne; DY1857) according to the manufacturer’s instructions. Frozen fecal pellets were weighed and resuspended at 100 mg/mL in PBS containing 0.1% Tween-20, vigorously homogenized by vortexing, and clarified by centrifugation at maximum speed for 1 min. The resulting supernatant was collected, stored at −80 °C, and subsequently used for ELISA analysis.

### Immunohistochemistry

Mice were anesthetized with an i.p. injection of 200 mg/kg pentobarbital and perfused transcardially with an ice-cold 1xPBS, followed by 4% paraformaldehyde (PFA) in 1xPBS. The brain, DRG and colon tissues were post-fixed for 4-6 hrs at 4°C and transferred to 30% sucrose in 1xPBS for 2 days. Tissues were frozen in O.C.T. (Tissue-Tek) and sectioned with a cryostat at 40-µm thickness. Sliced tissues were mounted on histology slides and immersed in blocking solution containing 5% normal donkey or goat serum and 0.3% Triton X-100 in 1xPBS for 1 hr at 4°C. The tissue slides were then incubated with primary antibodies at 4°C overnight. On the next day, tissues underwent extensive wash with clean 1xPBS followed by incubation with appropriate secondary antibody conjugated with AlexaFluor for 2 hrs at room temperature. After washing with 1xPBS for 5 min three times, stained slides were coverslipped with ProLong Diamond Antifade Mountant (Thermo Fisher, P36970). Histological images were acquired using either a Leica SP8 or Zeiss LSM 980 confocal microscope at a resolution of 1024 × 1024 pixels, using 10× or 20× air objectives with z-stack step sizes of 2–5 µm. Immunofluorescence signals were quantified using ImageJ (NIH, v1.53f). For each histological section, 8-10 consecutive z-slices were acquired and combined using an average-intensity z-projection. Projected images were background-subtracted and thresholded to generate binary masks of fluorescent signals. Automated particle/cell counting was performed to quantify the number of detected fluorescent cells. The following primary antibodies were used: anti-cFOS (Cell Signaling, #2250), anti-RFP (Rockland #48710), anti-Nav1.8 (Alamone Labs #ASC028), anti-CD45 (eBioscience #14-0451-82), anti-GFP (Thermo Fisher #A10262); secondary antibodies were used: Alexa Fluor^TM^ 555-conjugated donkey anti-rabbit (Invitrogen #A31572), Alexa Fluor^TM^ 594-conjugated goat anti-rabbit (Invitrogen #A11012), Alexa Fluor^TM^ 488-conjugated goat anti-mouse (Invitrogen #A11001), Alexa Fluor^TM^ 488-conjugated goat anti-chicken (Invitrogen #A11039).

### Fluorescence in situ hybridization

Mice underwent transcardial perfusion with 4% PFA and brains were post-fixed overnight and 30% sucrose for 48 hrs. Tissues were fresh-frozen in O.C.T. compound (Thermo Fisher Scientific) and stored at −80 °C until cryoslicing. Coronal sections of 20 µm thickness covering the entire PBN region were serially collected and mounted onto Superfrost Plus slides (Thermo Fisher Scientific) and stored at −80 °C for processing. RNAscope in situ hybridization was performed using the RNAscope Fluorescent Multiplex V2 Kit (Advanced Cell Diagnostics) following the manufacturer’s instructions. Slides were dehydrated through graded ethanol (50%, 70%, and 100%) and treated with hydrogen peroxide for 10 min, washed, and incubated with Protease IV for 30 min at room temperature. The following probes from Advanced Cell Diagnostics were used: Mm-Fos (316921), Mm-Nts (420441) and Mm-Crh (316091). Sections were incubated for probe hybridization and signal amplification steps in a HybEZ oven (ACD Bio) at 40 °C. Fluorescent signal detection was achieved using TSA Plus HRP reagents and Opal 520, 570, and 620 dyes (Akoya Biosciences). HRP reagents were applied for 15 min and Opal dyes for 30 min at 40 °C with 15min HRP blocking steps between each channel and wash buffer rinses between all incubations. FISH images covering serial sections of the PBN_L_ between bregma −5.0 mm and −5.4 mm were analyzed using ImageJ. Image brightness and contrast were adjusted using consistent thresholding parameters to optimize visualization of each fluorophore. Cellular boundaries were defined based on DAPI labeling, and individual mRNA puncta (5–10 pixels in diameter) were quantified within each cell. Cells containing >5 puncta of Fos, Nts, or Crh mRNA were classified as marker-positive. This threshold was applied consistently across all analyzed sections to determine the overlapping of gene-expressing populations.

### Statistical analysis

All statistical analyses were performed using GraphPad Prism. The specific statistical tests applied, exact p values, and sample sizes (n) are reported in the corresponding figures and figure legends. Data distribution was assessed where appropriate, and statistical tests were selected accordingly. Unless otherwise stated, all statistical tests were two-tailed, and statistical significance was defined as p < 0.05.

## Acknowledgments

We thank the members of the Woolf laboratory for inputs and discussions. We thank the IDDRC Animal Behavior and Physiology Core, funded by NIH/NICHD P50HD105351 and Cellular Imaging Core, funded by NIH P50 HD105351 and Viral Core, funded by NIH 5P30EY012196 at Boston Children’s Hospital; Dr. Mark L. Andermann for helpful comments and Dr. Fan Wang for general support. This study is supported by the National Institutes of Health grants R01AT011447 and R35NS105076 (C.J.W.), R01DK135707 and R01DK130836 (M.R), and the William Randolph Hearst Fund (Y.T.C). Schematics and illustrations were adapted from BioRender.com.

## Author Contributions

Y.T.C and C.J.W. conceived and designed the study. Y.T.C performed all the experiments and analyzed the data, with assistance from N.M.B. (AAV injections and histology), Y.J. (behavior scoring and RNAscope), J.R.R. (behavior scoring and histology), M.D.S. (GI transit assay and colon histopathology), P.E.M. (barostat recording), Y.L. (fiber photometry), O.B. (automatic behavior screening), K.H. (histology), B.Z. (behavior scoring) and B.L.T. (behavior assays). M.R. provided resources and oversight of the colon histopathology, GI transit and barostat recording experiments. Y.T.C and C.J.W. prepared the figures and wrote the manuscript with inputs from all the authors.

## Disclosures

C.J.W. is a founder of Nocion Therapeutics, Quralis, and BlackBox Bio, and serves on the scientific advisory boards of Lundbeck Pharma, Axonis, Niroda, Mimetic Medicine, Tenvie Therapeutics, and Tafalgie Therapeutics. M.R. collaborates with Fzata and her spouse is a Novo Nordisk employee.

## Notes

### Summary of Updates

Author Affiliations, Competing Interest Statement and Funder Information Declared updated.

