## Supplementary materials for "Colitis-induced visceral pain recruits central neurotensin neurons that modulate colonic sensitivity"

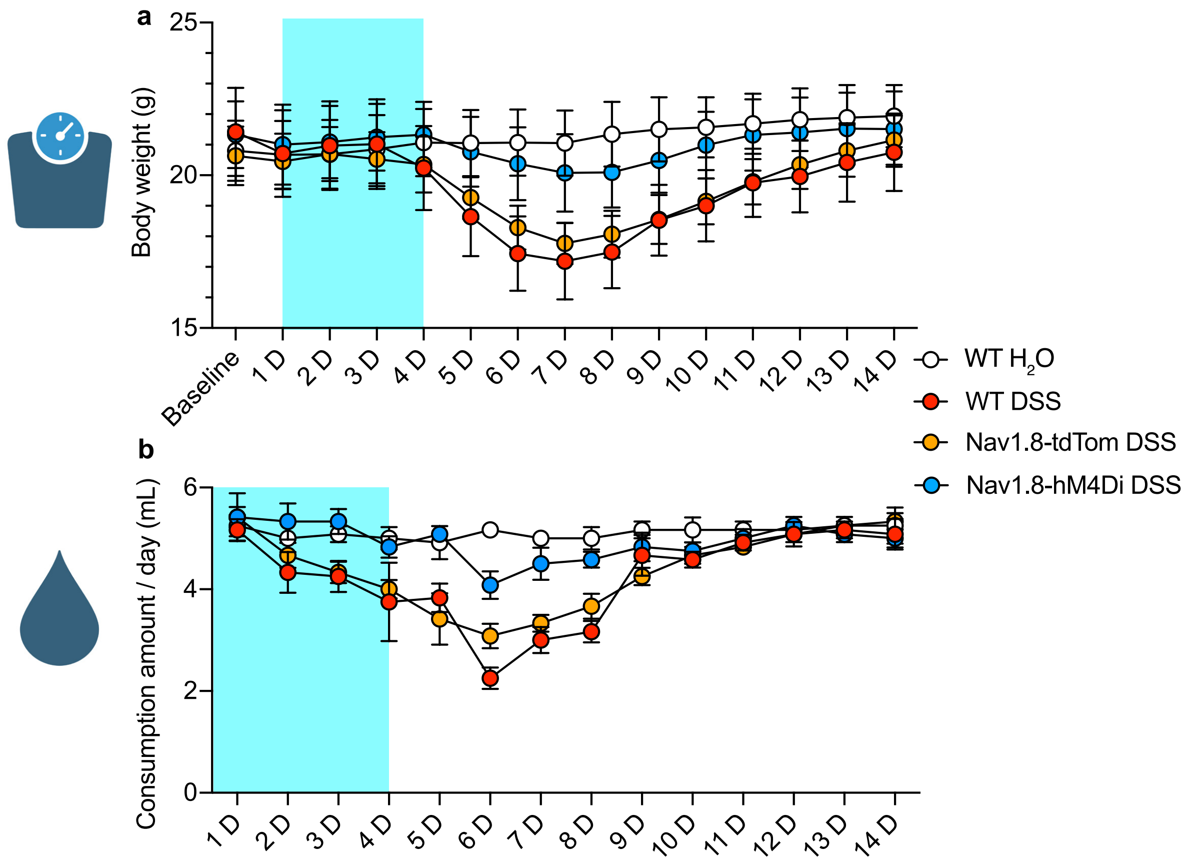


**Extended Fig 1. 4-day 3% DSS treatment leads to body weight loss and a reduction of drinking water intake that gradually recovers over 2 weeks.** (a) Measurement of body weight in WT mice with regular drinking water (n=6 mice), WT mice with 3% DSS drinking water (n=6 mice), Nav1.8-tdTom mice with 3% DSS drinking water (n=6 mice) and Nav1.8-hM4Di mice with 3% DSS drinking water (n=6 mice). In the two Nav1.8-Cre mice groups, 0.1 mg/kg of DCZ was injected i.p. at D1, D2, D3 and D4, once per day. (b) Measurement of water consumption in WT mice with regular drinking water (n=6), WT mice with 3% DSS drinking water (n=6), Nav1.8-tdTom mice with 3% DSS drinking water and DCZ exposure (n=6 mice) and Nav1.8-hM4Di mice with 3% DSS drinking water and DCZ (n=6 mice). Light blue shade: 3% DSS drinking water from D1-D4. Error bars represent s.e.m.

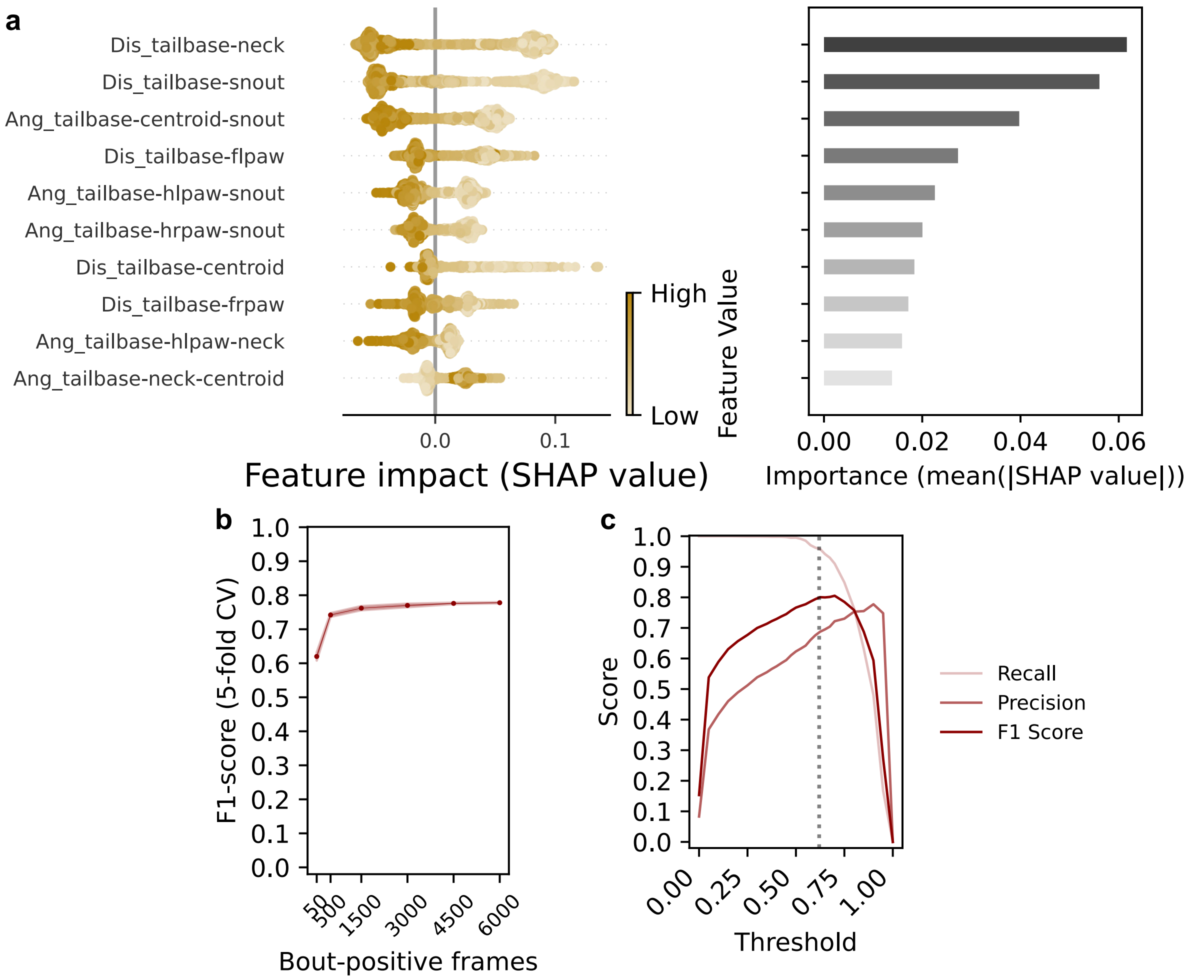


**Extended Fig 2. Evaluation of the algorithmic performance of licking behavior classifier.** (a) The top ten features for the licking classifier identified and interpreted using SHAP (Shapley Additive Explanations). The summary plot of the impact of each feature on the model's prediction: positive SHAP values indicate a higher probability of a bout-positive frame, while negative values contribute to a bout-negative classification (left). Each point represents an individual observation from a balanced subsample of 2,000 selected features. The gold scale reflects the scaled feature value. Horizontal bar graph of features ranked by their importance weights based on the mean absolute SHAP values (right). (b) F1-score learning curve for licking behavior classifier training. Data represents mean performance across 5-fold cross-validation at each training increment. Light shaded areas represent s.e.m. (c) Validation performance of the licking behavior classifier as a function of discrimination threshold. The curves represent F1-score (dark red), precision (red), and recall (light red). Gray dashed line denotes the optimal threshold value.

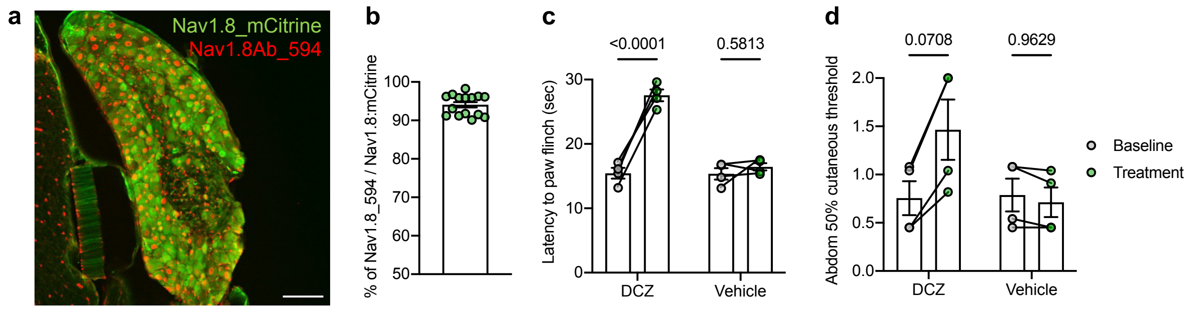


**Extended Fig 3. Histological and functional validation of Nav1.8-Cre mice.** (a) Representative histological image of an L6 dorsal root ganglion (DRG) from a Nav1.8-hM4Di/mCitrine mouse stained with an anti-Nav1.8 antibody conjugated with Alexa 594. Scale bar, 200 µm. (b) Percentage of signal overlap between Nav1.8-mCitrine-positive cells (green) and anti-Nav1.8 antibody staining (red). N=15 DRGs collected from L5-L6 segments of 3 mice. (c) Effect of Nav1.8-expressing nociceptor silencing on hind paw sensitivity towards noxious heat using a 52°C hotplate assay. DCZ n=4 mice, Vehicle n=4 mice. Two-way ANOVA with Šídák's multiple comparisons. (d) Effect of Nav1.8-expressing nociceptor silencing on mechanical sensitivity of the lower abdomen using an up-down von Frey assay. DCZ n=4 mice, Vehicle n=4 mice. Two-way ANOVA with Šídák's multiple comparisons.

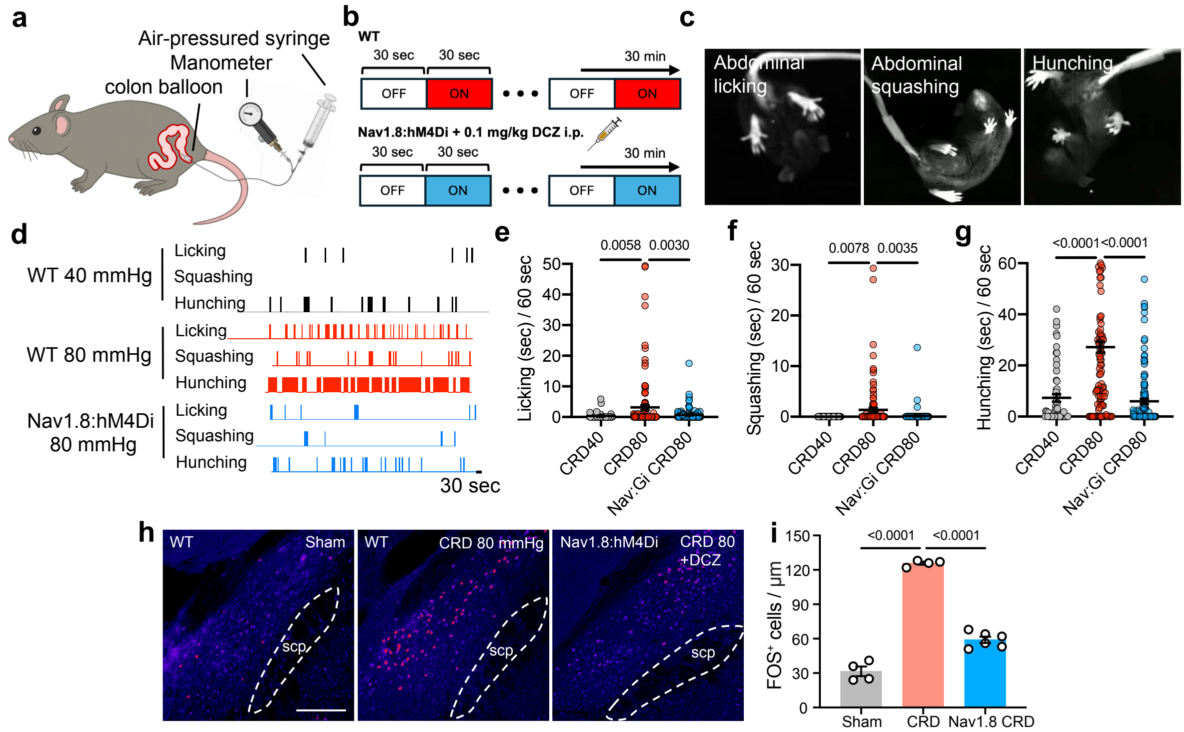


**Extended Fig 4. Nocifensive behaviors elicited by noxious colon distension.** (a) Illustration of a mouse with an intracolonic balloon insertion. A balloon connected to pressure-equalization tubing was inserted into the distal colon of mice with a manometer positioned at the bifurcation to monitor infusion pressure in real-time. (b) Experimental scheme to measure nocifensive behaviors in WT and Nav1.8-hM4Di DCZ-treated mice elicited in response to repeated colorectal distension (CRD) at 40 or 80 mmHg on for 30 secs then off for 30 seconds over a 30-min recording session. 0.1 mg/kg DCZ was injected i.p 15 min prior to the recording. (c) Example of mice showing abdominal licking (left), abdominal squashing (middle) and hunching (right) in a bottom-up recording platform. (d) Representative raster plots of mice exhibiting licking, squashing and hunching bouts in response to CRD at 40 and 80 mmHg of distension pressure, and after Nav1.8-lineage nociceptor silencing. (e, f, g) Time spent abdominal licking (e), abdominal squashing (f) and hunching (g) every 60-sec trial at CRD 40 (n=60 trials from 2 mice), CRD 80 (n=120 trials from 4 mice) and CRD 80 with Nav1.8-nociceptor silencing (Nav:Gi) (n=120 trials from 4 mice) over 30-min recording sessions. One-way ANOVA with Tukey's multiple comparisons. (h) Histological images of FOS staining in $PBN_L$ region in response to sham (left), CRD 80 mmHg (middle) and CRD 80 mmHg with Nav1.8-nociceptor silencing (right). scp, superior cerebellar peduncle. Scale bar, 200 µm. (i) Quantification of $FOS^+$ cells, sham n=2 mice, CRD 80 n=2 mice and Nav1.8-hM4Di + DCZ n=3 mice. One-way ANOVA with Tukey's multiple comparisons.

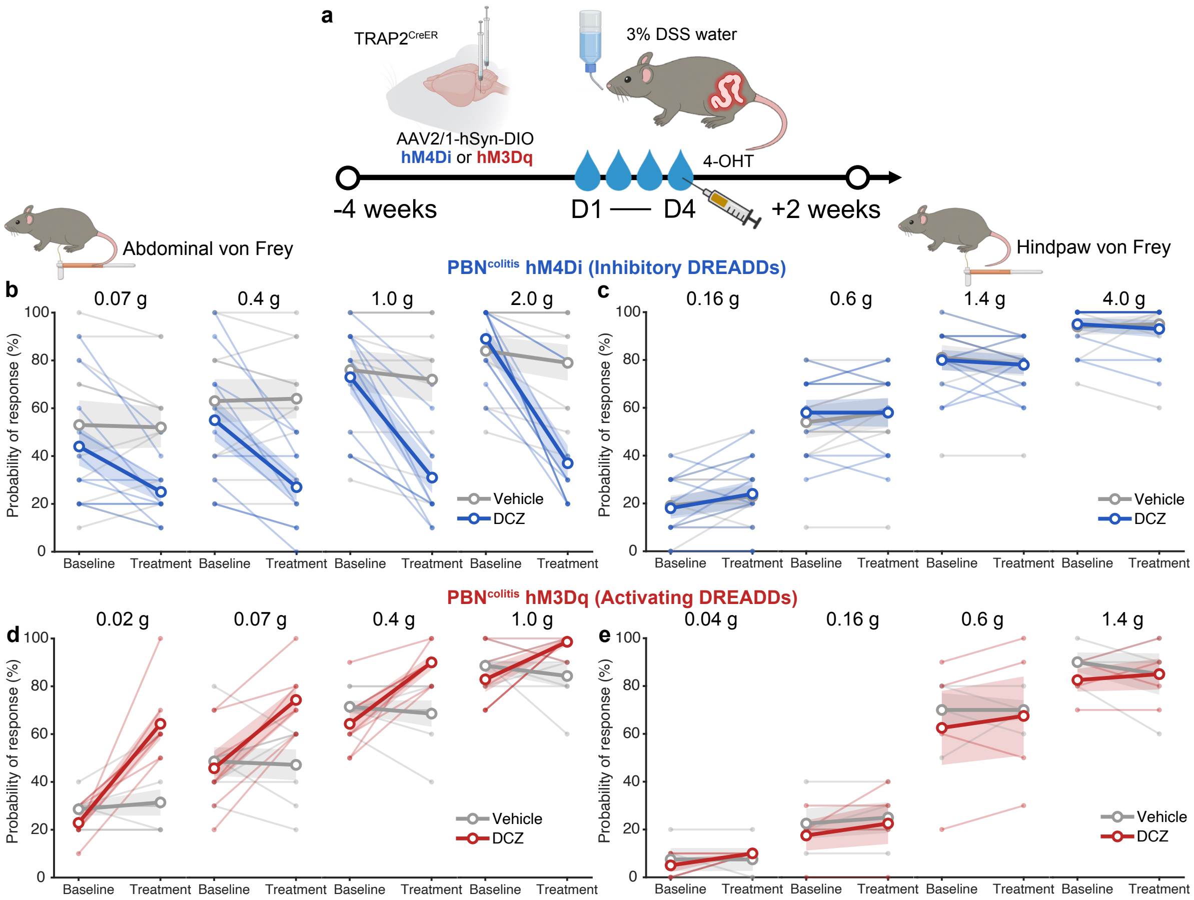


**Extended Fig 5. $\text{PBN}^{\text{colitis}}$ neurons modulate cutaneous sensitivity of the abdomen but have no effect on the hindpaw.** (a) Experimental strategy to functionally label DSS colitis-activated $\text{PBN}_\text{L}$ neurons. $\text{TRAP2}^{\text{CreER}}$ mice were injected with Cre-dependent AAV vectors encoding chemogenetic actuators (inhibitory hM4Di or activating hM3Dq) bilaterally into the $\text{PBN}_\text{L}$ region. 3% DSS drinking water was fed for 4 days followed by an i.p. injection of 4-hydroxytamoxifen (4-OHT) to drive expression of FOS-presenting cells. (b) Silencing effect of $\text{PBN}^{\text{colitis}}$ neurons using the inhibitory hM4Di, on abdominal sensitivity in response to a series of increased von Frey probing force at 0.07, 0.4, 1.0 and 2.0g (DCZ n=10 mice, Vehicle n=10 mice). (c) Silencing effect of $\text{PBN}^{\text{colitis}}$ neurons using an inhibitory hM4Di, on hindpaw sensitivity in response to von Frey forces of 0.16, 0.6, 1.4 and 4.0g (DCZ n=10 mice, Vehicle n=10 mice). (d) Effect of activating $\text{PBN}^{\text{colitis}}$ neurons using hM3Dq on abdominal sensitivity detected by von Frey probes of 0.02, 0.07, 0.4 and 1.0g (DCZ n=7 mice, Vehicle n=7 mice). (e) Activating effect of $\text{PBN}^{\text{colitis}}$ neurons using hM3Dq on hindpaw sensitivity in response to a series of increased von Frey probing force at 0.04, 0.16, 0.6 and 1.4g (DCZ n=4 mice, Vehicle n=4 mice). 0.1 mg/kg of DCZ was injected i.p 15 min prior to behavior test. Light shades represent s.e.m.

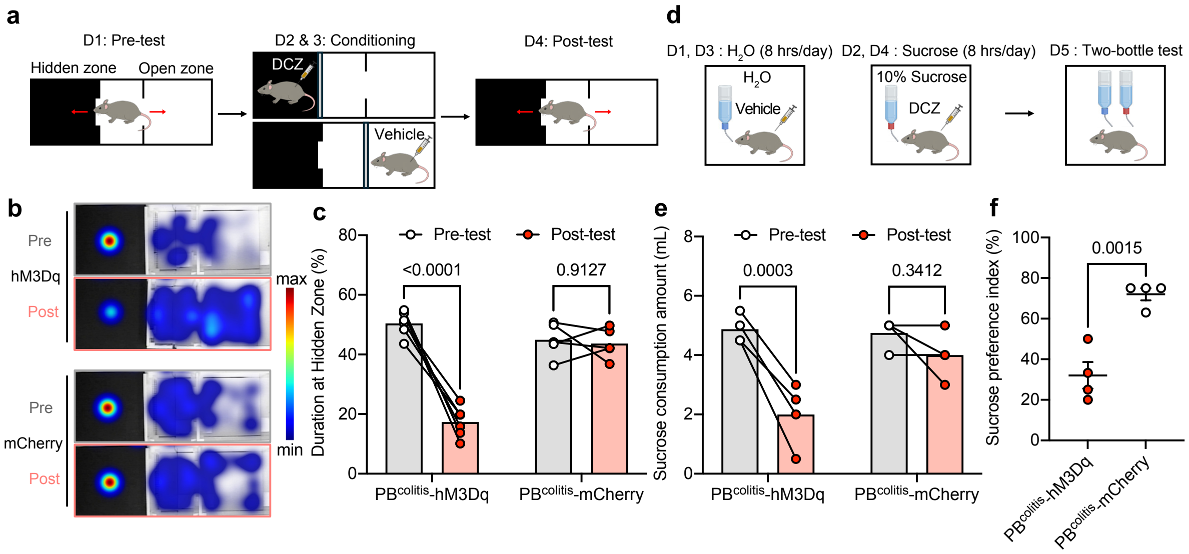


**Extended Fig 6. PBN$^{colitis}$ neurons drive aversive states.** (a) Experimental paradigm to measure affective valence of PBN$^{colitis}$ neurons on place motivation. Mice were placed in a three-chamber arena consisted of a baseline pre-test (D1), context-specific DCZ or vehicle-paired conditioning (D2-3), and a post-test (D4) to evaluate the shifted motivation towards the DCZ-paired chamber. (b) Representative occupancy heatmaps from pre-(grey) and post-tests (pink) of a conditioned place paradigm in mice expressing hM3Dq or mCherry in PBN$^{colitis}$ neurons. (c) Quantification of occupancy duration in the DCZ-paired hidden zone from pre- and post-tests in hM3Dq-expressing mice compared with mCherry controls (hM3Dq group n=6 mice, mCherry group n=5 mice, two-way ANOVA with Šídák's multiple comparisons). (d) Experimental paradigm to test the effect of PBN$^{colitis}$ neurons on taste motivation. Mice were exposed to water (D1, D3) or 10% sucrose (D2, D4) for 8 hours per day. Vehicle was administered on water days, whereas DCZ was administered during sucrose access. On D5, mice underwent a two-bottle choice test to measure intake preference. (e) Quantification of consumption of the DCZ-paired sucrose solution from pre- and post-tests in hM3Dq-expressing mice versus mCherry controls (hM3Dq group n=4 mice, mCherry group n=4 mice, two-way ANOVA with Šídák's multiple comparisons). (f) Two-bottle test (D5) to measure appetite preference in hM3Dq- and mCherry-expressing mice. Preference index = sucrose consumption / (sucrose + water consumption) x 100% (hM3Dq group n=4 mice, mCherry group n=4 mice, two-tailed unpaired t-test).

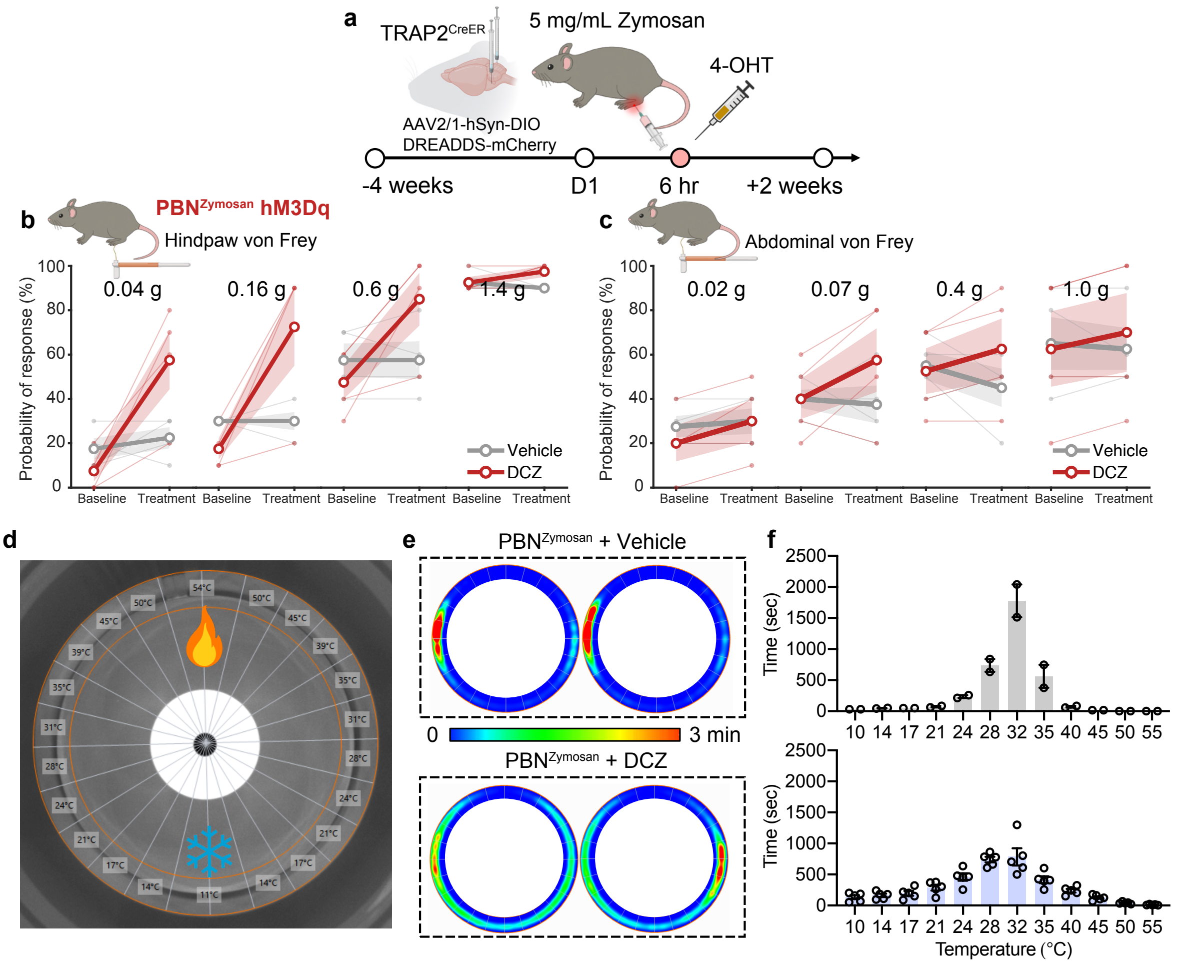


**Extended Fig 7. PBN$^{Zymosan}$ neurons tune hindpaw sensitivity but exert only a mild effect on abdominal sensitivity.** (a) Experimental strategy to gain functional access into the PBN$_L$ neurons in response to plantar zymosan-induced inflammatory pain (PBN$^{Zymosan}$) using TRAP2$^{CreER}$ mice with bilateral injection of Cre-dependent AAV vectors encoding inhibitory hM4Di or activating hM3Dq. 4-OHT was i.p. injected 6 hours following an intraplantar injection of 5mg/mL zymosan into the left hindpaw. (b) Activating effect of PBN$^{Zymosan}$ neurons using activating hM3Dq on cutaneous sensitivity of the hindpaw in response to a series of increased von Frey probing forces at 0.04, 0.16, 0.6 and 1.4g (DCZ n=4 mice Vehicle n=4 mice). (c) Effect of activating PBN$^{Zymosan}$ neurons using hM3Dq on cutaneous sensitivity of the abdomen in response to a series of increased von Frey probing forces at 0.02, 0.07, 0.4 and 1.0g (DCZ n=4 mice Vehicle n=4 mice). (d) Dorsal view of the thermal gradient ring (TGR) device to test the thermal sensitivity and preference over a temperature range of ~10°C-55°C. (e) Representative occupancy heat maps showing the outcome of silencing PBN$^{Zymosan}$ neurons with control vehicle (top) and DCZ (bottom) on occupancy duration across the TGR device over a 1-hour recording session. (f) Quantification of occupancy duration in each temperature zone for vehicle control (top n=2 mice) and after PBN$^{Zymosan}$ silencing with DCZ (bottom n=5 mice). 0.1 mg/kg of DCZ was injected i.p 15 min prior to behavior test. Light shades represent s.e.m.

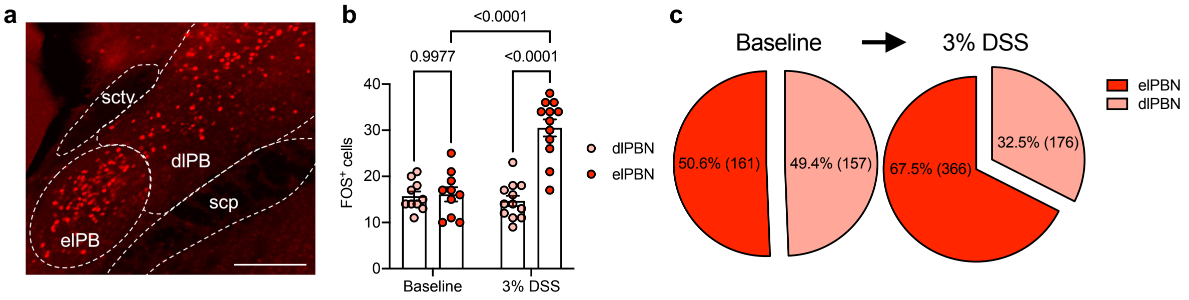


**Extended Fig 8. DSS-induced colitis preferentially activates the elPBN population.** (a) Representative image of a mouse PBN$_L$ with FOS antibody staining (red). Dashed lines outline subdivisions with the PBN$_L$ and neighboring fiber tracts. elPBN: external lateral parabrachial nucleus, dlPBN: dorsal lateral parabrachial nucleus, sctv: ventral spinocerebellar tract, scp: superior cerebellar peduncle. Scale bar, 200 μm. (b) Quantification of FOS$^+$ signals within elPBN and dlPBN at baseline (n=10 from 5 mice) and following 4-day 3% DSS treatment to induce colitis (n=12 from 6 mice). Two-way ANOVA with Tukey's multiple comparisons. (c) Pie chart summary of percentage of elPBN versus dlPBN at baseline (left) and DSS-induced colitis (right).

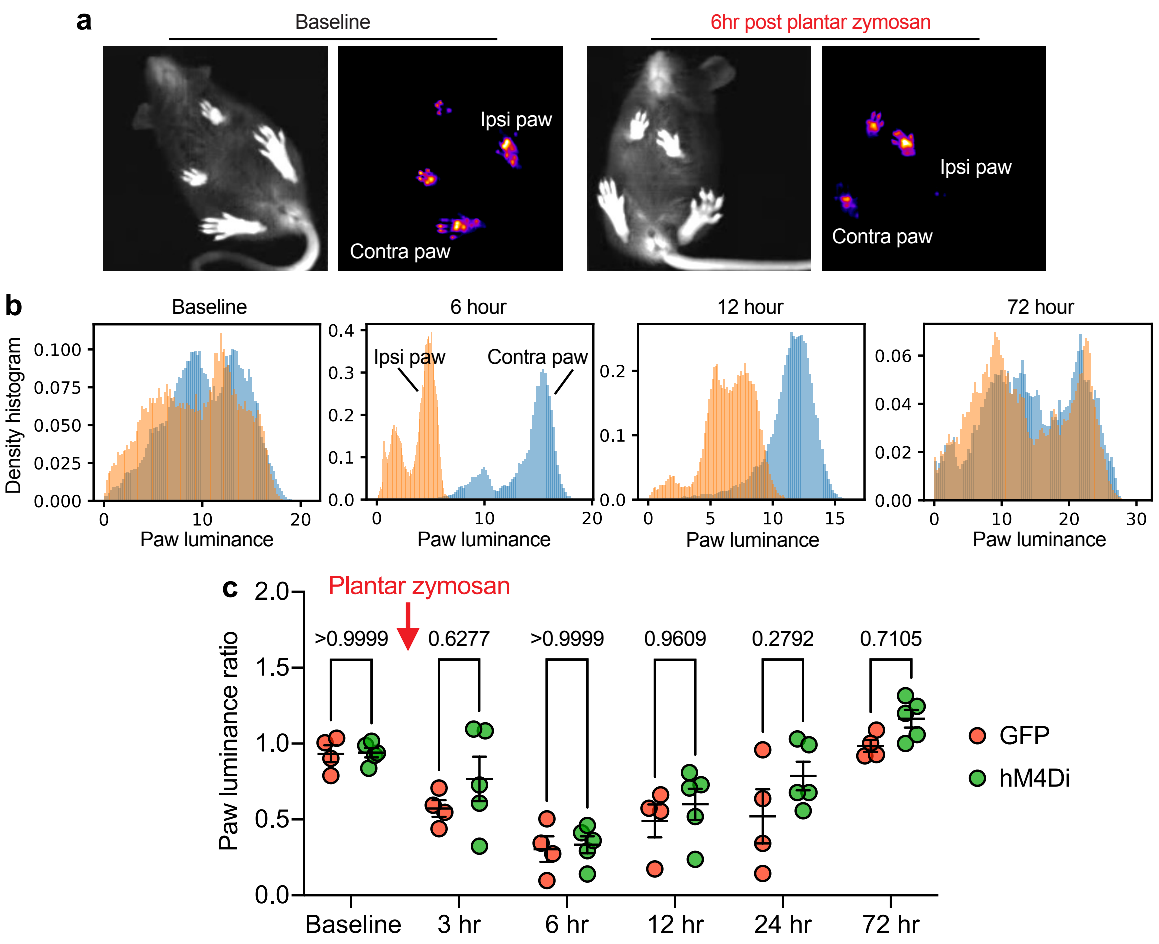


**Extended Fig 9. $\text{PBN}_\text{L}$ neurotensin neurons display limited modulatory effect on hindpaw hypersensitivity induced by zymosan injection.** (a) Representative behavior frames recorded from a bottom-up camera system to capture transilluminated signals (body frame, left) and frustrated total internal reflection (FTIR) signals (footprint, right) at baseline and post-zymosan treatment. (b) Representative density histograms of a mouse ipsilateral (orange) and contralateral (blue) paw luminance over a 30-minute recording at baseline and 6, 12, 72 hours after an intraplantar zymosan injection. (c) Paw luminance ratio calculated from NT-Cre mice received Cre-dependent AAVs encoding GFP or inhibitory hM4Di (GFP n=4 mice, hM4Di n=5 mice). 0.1 mg/kg of DCZ was injected i.p at 4-hour post-zymosan treatment in both groups. Paw luminance ratio = Luminance$^{\text{ipsilateral paw}}$/Luminance$^{\text{contralateral paw}}$. Two-way ANOVA with Šídák's multiple comparisons.

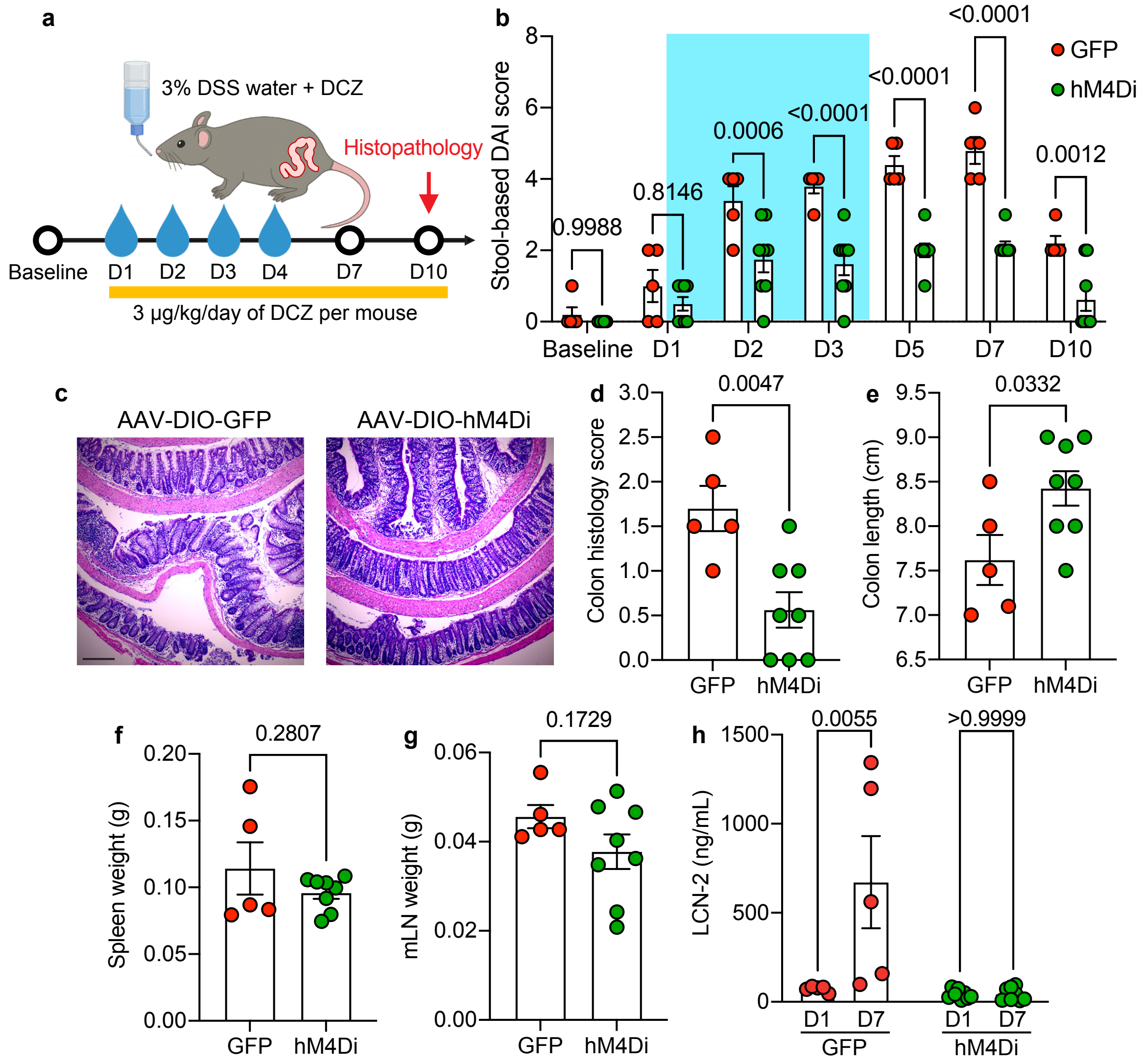


**Extended Fig 10. Prolonged silencing of $PBN_L$ neurotensin neurons improves colon pathology induced by colitis.** (a) Experimental scheme to evaluate the stool-based disease activity index (DAI) composed of scores for stool appearance and presence of blood in the stool using a Hemoccult card and histopathological assessment of tissues at D10. Concentration of 0.0286 mg/L DCZ was fed into the drinking water D1-D10 for mice to intake ~3 μg/kg of DCZ per day. (b) Quantification of stool-based DAI score in NT-Cre mice with bilateral injections of Cre-dependent AAV that encodes GFP (n=5 mice), or hM4Di (n=8 mice) under 3% DSS treatment for 4 days. Two-way ANOVA with Šídák's multiple comparisons. (c) Representative histology image of Hematoxylin and Eosin staining in the distal colon tissue from a mouse in the GFP control group (left) and mouse in the inhibitory hM4Di group (right). Scale bar, 200 μm. (d) Quantification of colon histology scores composed of crypt dropout (0-3), submucosal expansion (0-3) and neutrophil infiltrate (0-3). Quantification of colon length (e), spleen weight (f) and mesenteric lymph nodes (mLN) weight (g) in GFP (n=5 mice) and hM4Di group (n=8 mice). Two-tailed unpaired t-test. (h) Quantification of fecal lipocalin-2 levels (ng/mL) in GFP control (n=5 mice) and hM4Di group (n=8 mice) at 3% DSS water D1 and D7. Two-way ANOVA with Šídák's multiple comparisons.

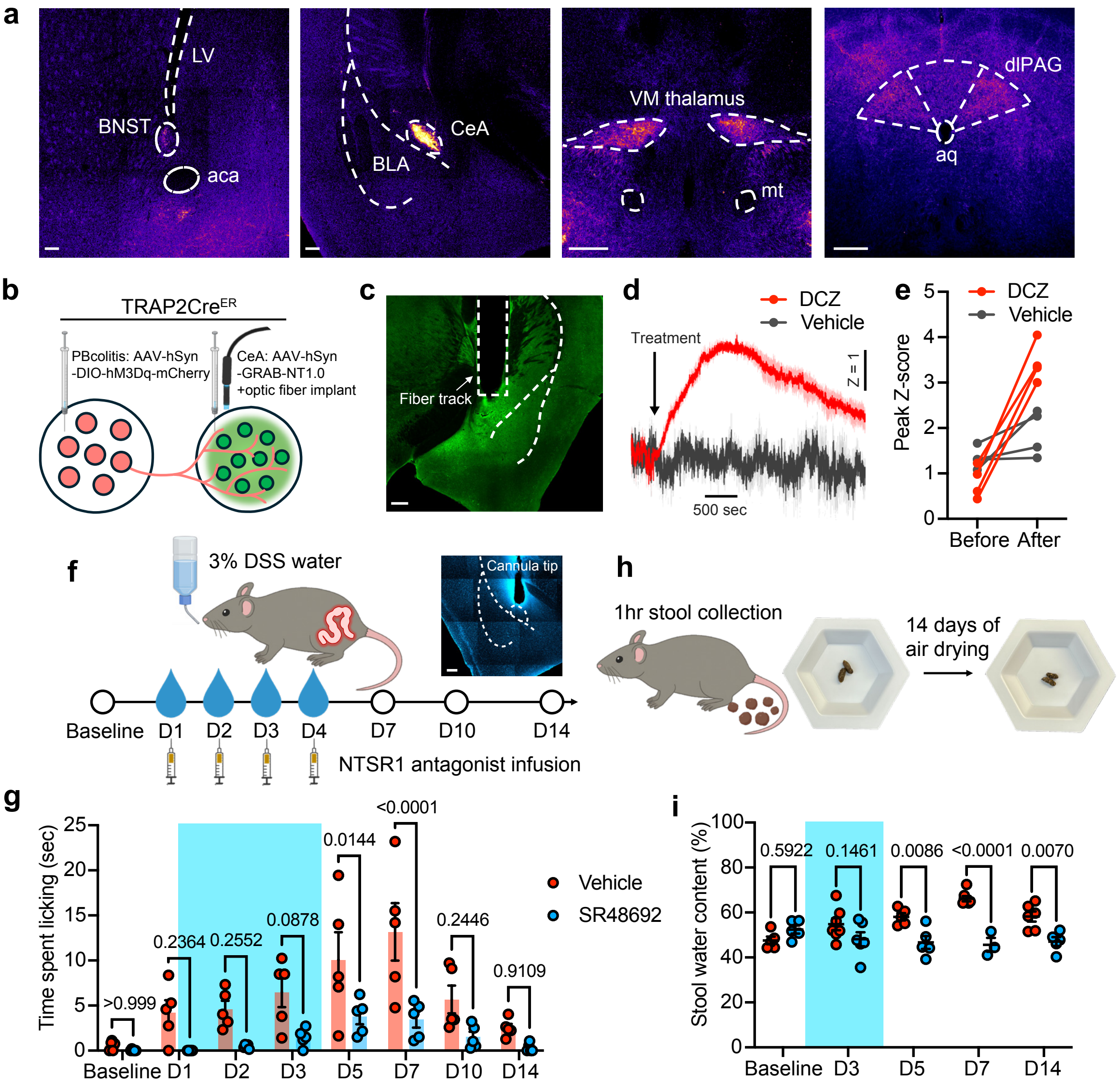


**Extended Fig 11. Targeting terminal neurotensin signaling ameliorates colitis-induced pain-related behavior.** (a) AAV-assisted projection mapping from $PBN_L$ NT neurons. Left to right: bed nucleus of the stria terminalis (BNST), lateral ventricle (LV), anterior commissure anterior part (aca), central amygdala (CeA), basolateral amygdala (BLA), ventral posteromedial thalamus (VM thalamus), mammillothalamic tract (mt), dorsolateral periaqueductal gray (dlPAG) and aqueduct (aq). Scale bars, 200 μm. (b) Schematic of functional connectivity of $PBN_L$-to-CeA circuit. Cre-dependent AAV encoding an activating DREADD (hM3Dq) was delivered into the $PBN^{colitis}$ and AAV-GRAB (GPCR-activation-based) sensor encoding neurotensin activity was injected into the CeA region followed by an optic fiber implant to measure terminal NT signaling *in vivo*. (c) Histological verification of AAV-hSyn-GRAB-NT1.0 injection and fiber implant track above the CeA region. Scale bars, 300 μm. (d) Average z-scored ΔF/F of spontaneous NT release in the CeA on chemogenetic activation of $PBN^{colitis}$ with an i.p. injection of DCZ (red, n=4 mice) versus vehicle control (grey, n=4 mice). Light shaded areas represent s.e.m. (e) Quantification of peak z-scored ΔF/F over 30 min before and after the drug treatment from (d). (f) Experimental scheme to evaluate therapeutic effect of SR48692, a selective NTSR1 antagonist, at an infusion dose of 20 nM, 0.3 µL (once daily D1-D4) into the CeA region of WT mice with 3% DSS drinking water. Inset: histological verification of cannula implant track above CeA. Scale bars, 200 µm. (g) Quantification of colitis-induced licking behavior using a supervised machine-learning classifier in WT mice with intracranial infusions of SR48692 (n=5 mice) or vehicle (n=5 mice) with 4-day 3% DSS treatment. Two-way ANOVA with Šídák's multiple comparisons. (h) Schematic of diarrhea evaluation by measuring stool weight following two weeks of air drying to calculate percentage of water content. (i) Quantification of stool water content in WT mice with infusion of SR48692 (Baseline n=5, D3 n=6, D5 n=5, D7 n=3, D14 n=5 mice) or vehicle (Baseline n=5, D3 n=7, D5 n=5, D7 n=7, D14 n=6 mice) under 4-day 3% DSS treatment. Two-way ANOVA with Šídák's multiple comparisons.
